# Specific *N*-glycans regulate an extracellular adhesion complex during somatosensory dendrite patterning

**DOI:** 10.1101/2021.10.11.464022

**Authors:** Maisha Rahman, Nelson J. Ramirez-Suarez, Carlos A. Diaz-Balzac, Hannes E. Bülow

**Author notes:** Institute of Science and Technology Austria, Am Campus 1, 3400, Klosterneuburg, Austria (Nelson J. Ramirez-Suarez). University of Rochester, Rochester, New York, 14627 (Carlos A. Diaz-Balzac).

## Abstract

*N*-glycans are molecularly diverse sugars borne by over 70% of proteins transiting the secretory pathway and have been implicated in protein folding, stability, and localization. Mutations in genes important for *N*-glycosylation result in congenital disorders of glycosylation that are often associated with intellectual disability. Here, we show that structurally distinct *N*-glycans regulate the activity of an extracellular protein complex involved in patterning of somatosensory dendrites in *Caenorhabditis elegans*. Specifically, *aman-2/Golgi alpha-mannosidase II*, a conserved key enzyme in the biosynthesis of specific *N*-glycans regulates the activity of the Menorin adhesion complex without obviously affecting protein stability and localization of its components. AMAN-2 functions cell-autonomously to ensure decoration of the neuronal transmembrane receptor DMA-1/LRR-TM with high-mannose/hybrid *N*-glycans. Moreover, distinct types of *N*-glycans on specific *N*-glycosylation sites regulate the DMA-1/LRR-TM receptor, which together with three other extracellular proteins forms the Menorin adhesion complex. In summary, specific *N*-glycan structures regulate dendrite patterning by coordinating the activity of an extracellular adhesion complex suggesting that the molecular diversity of *N*-glycans can contribute to developmental specificity in the nervous system.

## INTRODUCTION

Development of a nervous system in metazoans requires the coordinated interactions of extracellular molecules to ensure correct neuronal morphogenesis, and to establish connectivity (Jan & Jan, 2010; Dong *et al*, 2015; Lefebvre, 2021). Most of these extracellular proteins are glycoconjugates, i.e. carry different types of glycans attached to the protein backbone. Glycans are molecularly the most diverse molecules in nature, in part because they are not genetically encoded. Yet their structures are not random and are therefore conceptually attractive to broaden the molecular diversity and specificity of extracellular proteins and their interactions during development. For example, glycosaminoglycans, a class of glycans, have been suggested to modulate protein-protein interactions and provide information during development by way of their structural diversity (Holt & Dickson, 2005; Bülow & Hobert, 2006; Poulain & Yost, 2015; Masu, 2016). Whether the structural diversity of other classes of glycans such as *N*-glycans and *O-*glycans serve similar functions is unclear.

*N*-glycans are a structurally diverse group of glycans that fall into four classes: high-mannose, hybrid, complex, and paucimannose-type *N*-glycans, and are invariantly attached via an asparagine to a protein backbone (Stanley *et al*, 2015). Importantly, 70% of all proteins transiting the endoplasmic reticulum are post-translationally *N*-glycosylated (Apweiler *et al*, 1999). Therefore, the structural diversity of *N*-glycans could significantly expand the repertoire and specificity of protein interactions in the extracellular space. *N*-glycans in general have been shown to be important for protein folding, stability, and localization (Stanley *et al*., 2015). Moreover, mutations in genes involved in *N*-glycosylation in humans result in Congenital disorders of glycosylation (CDG), which are multi-syndromic and often include neurological symptoms, including intellectual disability (Freeze, 2006; Jaeken & Peanne, 2017; Chang *et al*, 2018; Ng & Freeze, 2018). Studies in vertebrates and invertebrates have shown that mutants that compromise *N*-glycan biosynthesis or *N*-glycan attachment result in defects in cell surface localization of cell adhesion molecules and axon guidance cues (Sekine *et al*, 2013; Medina-Cano *et al*, 2018; Mire *et al*, 2018). The question of whether and how specific classes of *N*-glycans modulate extracellular pathways or complexes during nervous system development has not been explored.

Here we use PVD somatosensory dendrites, which display complex and stereotyped branching patterns in the nematode *Caenorhabditis elegans* (Fig.1A) (Oren-Suissa *et al*, 2010; Smith *et al*, 2010; Albeg *et al*, 2011) to investigate the role of different classes of *N*-glycans during development. We found that *aman-2/Golgi alpha-mannosidase* II, a conserved enzyme important for the synthesis of complex and paucimannose-type *N*-glycans is required for PVD dendrite morphogenesis. Specifically, *aman-2/Golgi alpha-mannosidase II* ensures the correct decoration of the leucine-rich transmembrane receptor DMA-1/LRR-TM in PVD with high-mannose and paucimannose-type *N*-glycans on specific *N*-glycosylation sites. Rather than controlling trafficking or surface localization of DMA-1/LRR-TM, we provide evidence that correct *N*-glycosylation of DMA-1/LRR-TM is essential for the function of DMA-1/LRR-TM as part of the Menorin pathway during PVD patterning. This pathway comprises two conserved cell adhesion molecules SAX-7/L1CAM and MNR-1/Menorin that function from the epidermis, and a secreted chemokine LECT-2/Chondromodulin II from muscle that together with DMA-1/LRR-TM form a high affinity cell adhesion complex (Fig.1B) (Inberg *et al*, 2019; Sundararajan *et al*, 2019). Together, our experiments suggest that distinct classes of *N*-glycans serve specific functions beyond protein folding and localization, and can contribute to developmental specificity during neuronal morphogenesis.

**Figure 1.**
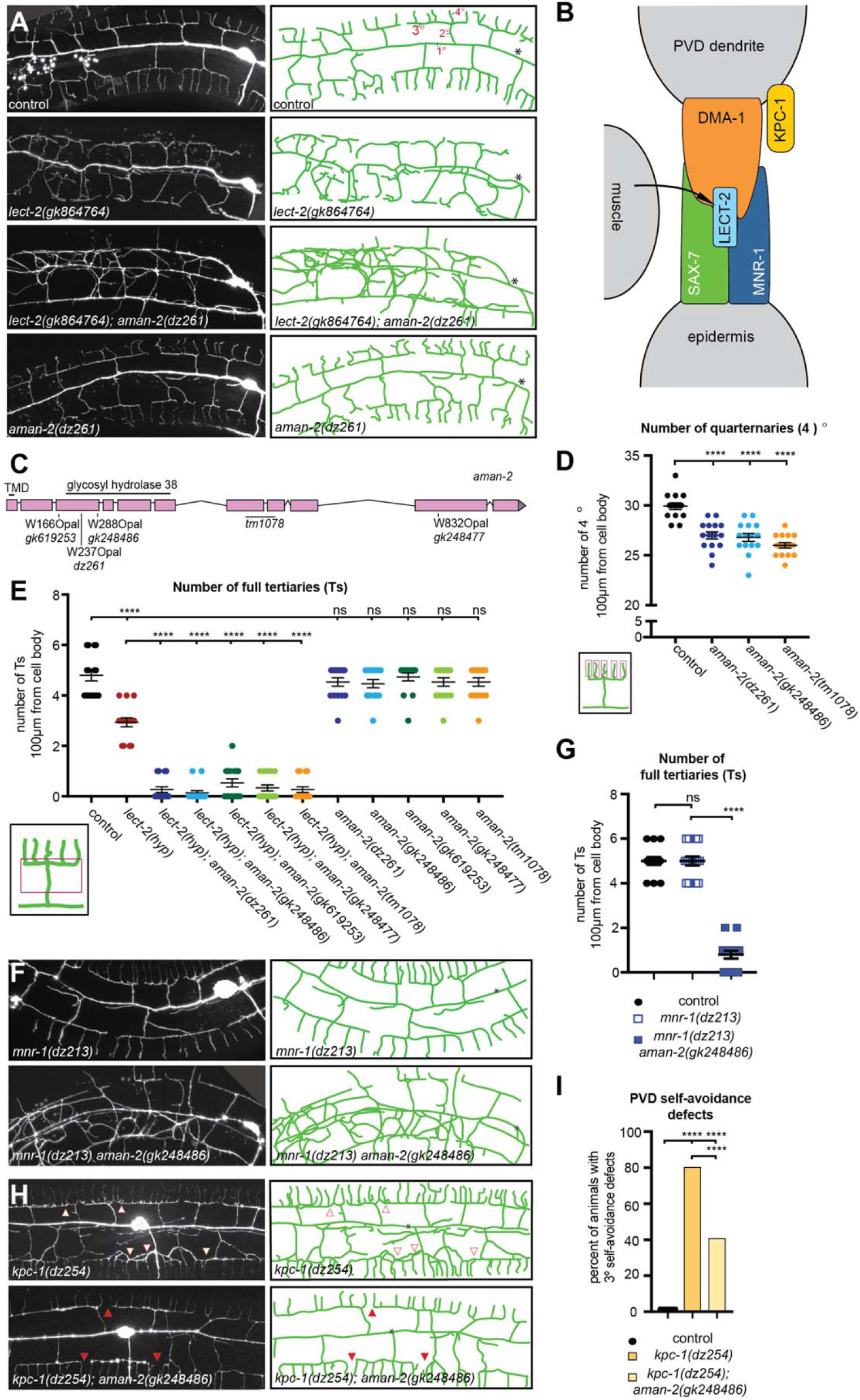
AMAN-2/Golgi alpha-mannosidase II is required for PVD dendrite patterning. (A) Fluorescent images (left panels) and tracings (right panels) of PVD of the indicated genotypes. PVD is visualized by the *wdIs52* transgene. Primary (1°), secondary (2°), tertiary (3°), and quaternary (4°) dendrites are indicated, and the cell body is marked with an asterisk. Anterior is to the left and dorsal is up in all panels. (B) Schematic of the Menorin complex, including DMA-1/LRR-TM, SAX-7/L1CAM, MNR-1/Menorin, and LECT-2/Chondromodulin II as well as the negative regulator KPC-1/Furin. (C) Genomic environs of *aman-2*. An *N*-terminal transmembrane domain (TMD) is indicated as it encodes a type II transmembrane protein. All alpha-mannosidase II proteins contain a glycosyl hydrolase 38 domain. Four nonsense alleles, and one deletion allele (*tm1078*) of *aman-2* are denoted. (D) Quantification of the number of quaternary branches (indicated in schematic) in different *aman-2* alleles and in wild type control animals. All loss of function alleles of *aman-2* show a significant decrease in quaternary branch number. Data are represented as mean ± SEM. Statistical significance was calculated using the Kruskal-Wallis test and is indicated (****p ≤ 0.0001). n = 15 for all genotypes. (E) Quantification of the number of “Ts” (formed by secondary and tertiary branches as shown in schematic) in the genotypes indicated. Data are represented as mean ± SEM. Statistical significance was calculated using the Kruskal-Wallis test and is indicated (****p ≤ 0.0001; ns= not significant). n = 15 for all genotypes. (F) Fluorescent images and tracings of PVD in *mnr-1*(*dz213*) hypomorphic animals alone and in combination with an *aman-2*(*null*) mutant. The *dz213* introduces a L135F missense mutation in the DUF2181 domain of MNR-1/Menorin. PVD is visualized by the *dzIs53* transgene. The cell body is marked with an asterisk. (G) Quantification of the number of “Ts” in genotypes indicated and traced in (F). The loss of *aman-2* severely enhances the *mnr-1*(*dz213*) phenotype. Data are represented as mean ± SEM. Statistical significance was calculated using the Mann-Whitney test and is indicated (****p ≤ 0.0001; ns= not significant). n >23 for all genotypes. (H) Fluorescent images and tracings of PVD in *kpc-1*(*dz254*) hypomorph animals alone and in combination with an *aman-2*(*null*) mutant. PVD is visualized by the *dzIs53* transgene. White arrows indicate self-avoidance defects in tertiary branches. Red arrows show gaps between tertiary branches (no self-avoidance defects). The cell body is marked with an asterisk. (I) Quantification of the percent of self-avoidance defects in genotypes indicated and traced in (G). The loss of *aman-2* suppresses defects in the *kpc-1*(*dz254*) phenotype. Data are represented as mean. Statistical significance was calculated using the Z-test and is indicated (****p ≤ 0.0001). n >15 for all genotypes.

## RESULTS

### The *N*-glycosylation enzyme AMAN-2/Golgi alpha-mannosidase II is required for PVD dendrite patterning

To identify additional factors that regulate the Menorin pathway, we performed an unbiased genetic screen for factors that modify a partial loss of function allele of the chemokine *lect-2/Chondromodulin II* (Diaz-Balzac *et al*, 2016; Zou *et al*, 2016). We isolated *dz261* as a strong enhancer of the partial loss of function *lect-2* mutation in addition to other alleles in known genes of the Menorin pathway (Fig.1A,D, Fig.EV1A-C). Using a combination of mapping, sequencing and transformation rescue we identified *dz261* as an allele of *aman-2/Golgi alpha-mannosidase II* (Fig.1C, Fig.EV1A-C), which encodes a central enzyme in the *N*-glycosylation biosynthetic pathway not previously implicated in dendrite development. The allele *dz261* is likely a complete loss of function mutation as it introduces an early stop codon (Fig.1C, Fig. EV1A-C). Quantifications of PVD branching patterns in *aman-2(dz261)* null mutants, as well as four additional nonsense/deletion alleles of *aman-2*, demonstrate that this Golgi alpha-mannosidase II is required for the formation of quaternary branches, but not for the formation of secondary or tertiary branches of PVD dendrites (Fig.1D,E, Fig. EV1D). These observations are reminiscent of hypomorphic alleles of *dma-1/LRR-TM* (Tang *et al*, 2019) and suggest that *aman-2* may be necessary for full functionality of the Menorin pathway.

### AMAN-2/Golgi alpha-mannosidase II positively regulates the Menorin pathway

To directly test the genetic relationship between *aman-2/Golgi alpha-mannosidase II* and the Menorin pathway, we performed double mutant analyses. We found that the loss of *aman-2/Golgi alpha-mannosidase II* strongly enhances the severity of PVD branching defects in partial loss-of-function mutants of *lect-2/Chondromodulin II* (*gk864764*) and *mnr-1/Menorin* (*dz213*, Ramirez-Suarez & Bülow, unpublished) (Fig.1A,E,F,G), both of which are essential components of the conserved Menorin cell adhesion complex, and act as positive regulators of PVD development (Fig.1E-G) (Dong *et al*, 2013; Salzberg *et al*, 2013; Diaz-Balzac *et al*., 2016; Zou *et al*., 2016). In contrast, loss of *aman-2/Golgi alpha-mannosidase II* suppressed the self-avoidance defects of tertiary dendrites in partial loss of a function mutation of *kpc-1/Furin* (*dz254*) (Fig.1H,I, Fig. EV1C), a known negative regulator of the Menorin pathway (Schroeder *et al*, 2013; Salzberg *et al*, 2014; Dong *et al*, 2016). Therefore, we conclude that *aman-2/Golgi alpha-mannosidase II* normally functions to positively regulate the Menorin pathway to ensure correct PVD dendrite patterning.

### *AMAN-2/Golgi alpha-mannosidase II* does not serve obvious functions in regulating transport or abundance of the DMA-1/LRR-TM

Mutations in *N*-glycosylation are often associated with protein folding defects and trafficking blocks due to misfolding (Stanley *et al*., 2015) and can, for example, result in lower abundance of cell surface proteins such as cell adhesion proteins in the nervous system (Medina-Cano *et al*., 2018). Because protein misfolding is more likely to occur at elevated temperatures (Gasser *et al*, 2008; Vabulas *et al*, 2010), we tested whether PVD branching defects in *aman-2/Golgi alpha-mannosidase II* mutations get progressively more severe with increasing temperatures. We found no significant increase in dendrite branching defects at 25°C compared to 15°C in *aman-2*(*gk248486*) mutant animals, in contrast to hypomorphic *lect-2*(*gk864764*) mutant animals (Fig. EV2A, B). Previous work showed that mutations causing a secretory block as a result of a defective unfolded protein response trap a DMA-1::GFP reporter in the cell body of PVD (Wei *et al*, 2015; Salzberg *et al*, 2017). We therefore questioned whether the loss of *aman-2/Golgi alpha-mannosidase II* can lead to defects in protein folding and trafficking, and a possible secretory block. We analyzed the amount and number of puncta of the DMA-1::GFP reporter in the soma, as well as in dendrite branches, and found that DMA-1::GFP fluorescence in both the soma and primary dendrites, and the number of DMA-1::GFP puncta in tertiary dendrites remained unaffected in *aman-2*(*gk248486*) mutant animals (Fig.EV2C-E). Moreover, localization or abundance of LECT-2/Chondromodulin II and SAX-7/L1CAM were also not obviously affected by loss of *aman-2/Golgi alpha-mannosidase II* (data not shown). Taken together, these findings suggest that AMAN-2/Golgi alpha-mannosidase II does not primarily function to ensure protein folding, stability, or transport of factors of the Menorin pathway, but may rather regulate more specific aspects of the Menorin pathway during PVD patterning.

### Enzymatic activity of AMAN-2/Golgi alpha-mannosidase II is required cell-autonomously in PVD to form higher order branches

The octasaccharide GnMan5Gn2 in a specific linkage configuration is the unique precursor to hybrid, complex, and paucimannose *N*-glycans (Fig.2A,B;Fig.3A) (Moremen, 2002; Paschinger *et al*, 2019). AMAN-2 is a Golgi alpha-mannosidase II, which is conserved from yeast to humans, and cleaves two specific mannose residues from GnMan5Gn2, thereby generating the substrate for formation of complex and paucimannose., *N*-glycans (Fig.2A,B;Fig.3A) (Moremen, 2002; Paschinger *et al*., 2019). To determine where AMAN-2/Golgi alpha-mannosidase II functions and whether enzymatic activity is required for its role in PVD dendrite morphogenesis, we investigated transgenic expression of a wildtype AMAN-2 cDNA under the control of heterologous promoters in PVD, muscle or epidermis for their ability to rescue *aman-2* mutant defects. Expression in PVD, but not muscle or epidermis rescued the enhanced phenotypes in PVD dendrite branching of *lect-2*(*gk864764*); *aman-2*(*gk248486*) and *mnr-1*(*dz213*) *aman-2*(*gk248486*) double mutants (Fig.2C). These observations suggest that AMAN-2/Golgi alpha-mannosidase II functions cell-autonomously to pattern PVD dendritic arbors.

**Figure 2.**
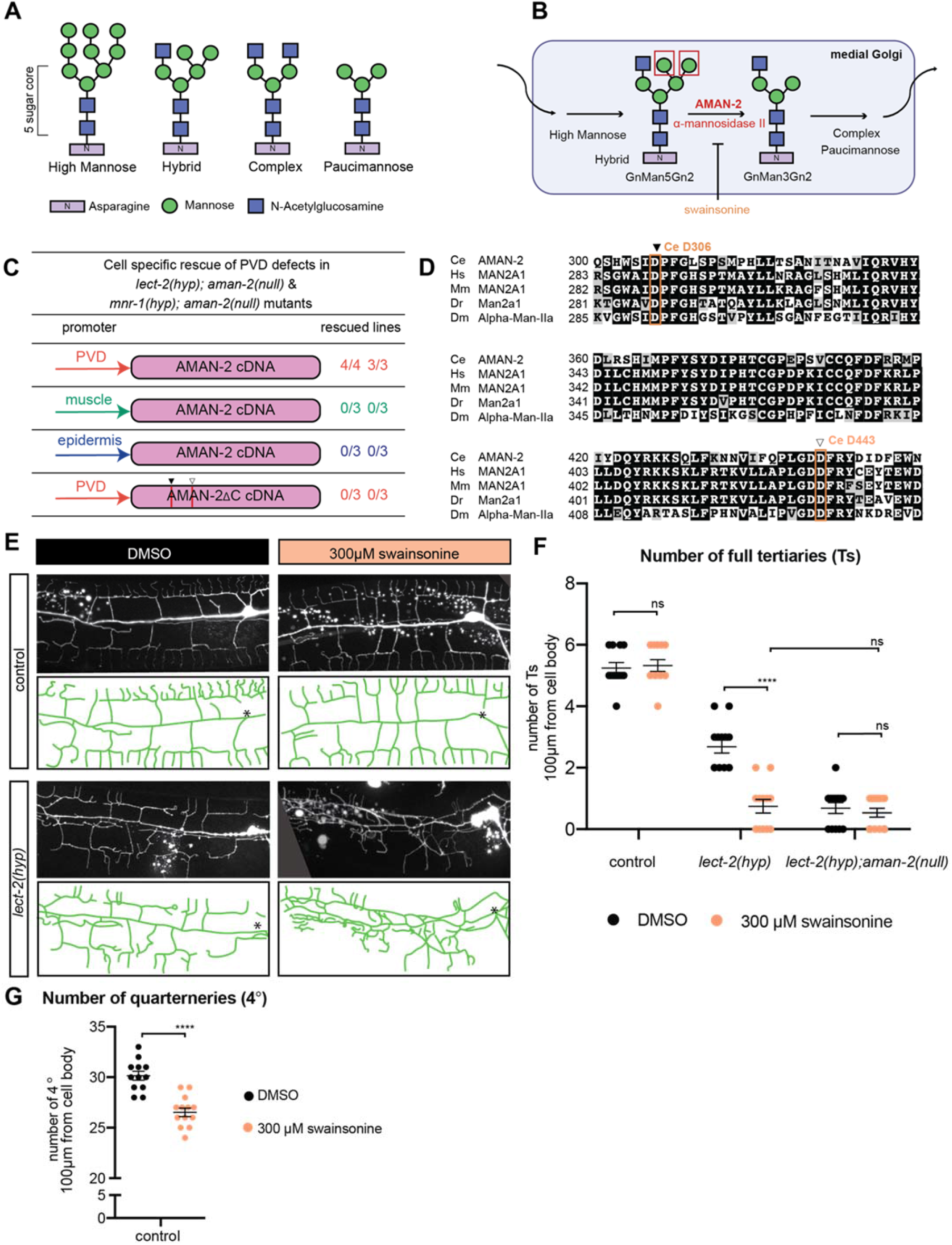
AMAN-2/Golgi alpha-mannosidase II requires enzymatic activity in PVD to form higher order branches. (A) Types of *N*-linked glycans. The shared penta-saccharide core consists of two *N*-acetylglucosamines (blue squares) and 3 mannoses (green circles) attached onto an Asparagine residue with an N-X-S/T consensus site. Glycan types vary by identity of additional sugars onto the pentasaccharide core. (B) Drawing showing the enzymatic activity of AMAN-2 in the medial Golgi. AMAN-2 cleaves two specific α1,3 and α1,6 mannose linked residues boxed in red, allowing for the formation of complex and paucimannose type *N*-glycans. Arrows denote other enzymes. Swainsonine specifically inhibits enzymatic activity of alpha-mannosidase II. (Man= mannose, Gn= *N*-acetylglucosamine). (C) Table showing cell-specific rescue experiments of PVD defects. AMAN-2 cDNA is expressed under the control of PVD, muscle, and epidermal specific promoters in the indicated double mutant backgrounds. Rescue is defined by restoration of the enhanced PVD phenotype back to that of the single hypomorphic mutants alone. 25 transgenic animals and their non-transgenic siblings were scored for each line. (D) Multiple sequence alignment of human MAN2A1 (Hs; acc# NP_002363.2), mouse MAN2A1 (Mm; acc# NP_032575.2), zebrafish Man2a1 (Dr; acc# NP_001103497.2), fruit fly Alpha-Man-IIa (Dm; acc# NP_650494.2) and *C. elegans* AMAN-2 (Ce; acc# NP_505995.2) created by COBALT (constrained based multiple sequence alignment tool). Conserved catalytic sites D306 (black arrow) and D443 (white arrow) are boxed in orange. (E) Fluorescent images and tracings of wild type control (top) and *lect-2(gk846764*) hypomorphic animals (bottom) fed on plates with 300μm swainsonine vs a DMSO control. PVD is visualized by the *wyIs581* transgene. The cell body is denoted with an asterisk. Anterior is to the left and dorsal is up in all panels. (F) Quantification of the number of “Ts” in denoted genetic backgrounds (*aman-2* null is *gk248486*). Black data points indicate DMSO and orange data points show swainsonine treated animals. Data are represented as mean ± SEM. Statistical significance was calculated using the Mann-Whitney test and is indicated (****p ≤ 0.0001; ns= not significant). n = 12 for all genotypes. (G) Quantification of the number of quaternary dendrites in wild type control animals fed on plates with and without 300μm swainsonine. Animals treated with swainsonine show a significant decrease in quaternary branch number, akin to the data in Figure 1C. Data are represented as mean ± SEM. Statistical significance was calculated using the Mann-Whitney test and is indicated (****p ≤ 0.0001). n = 12 for each experiment.

**Figure 3.**
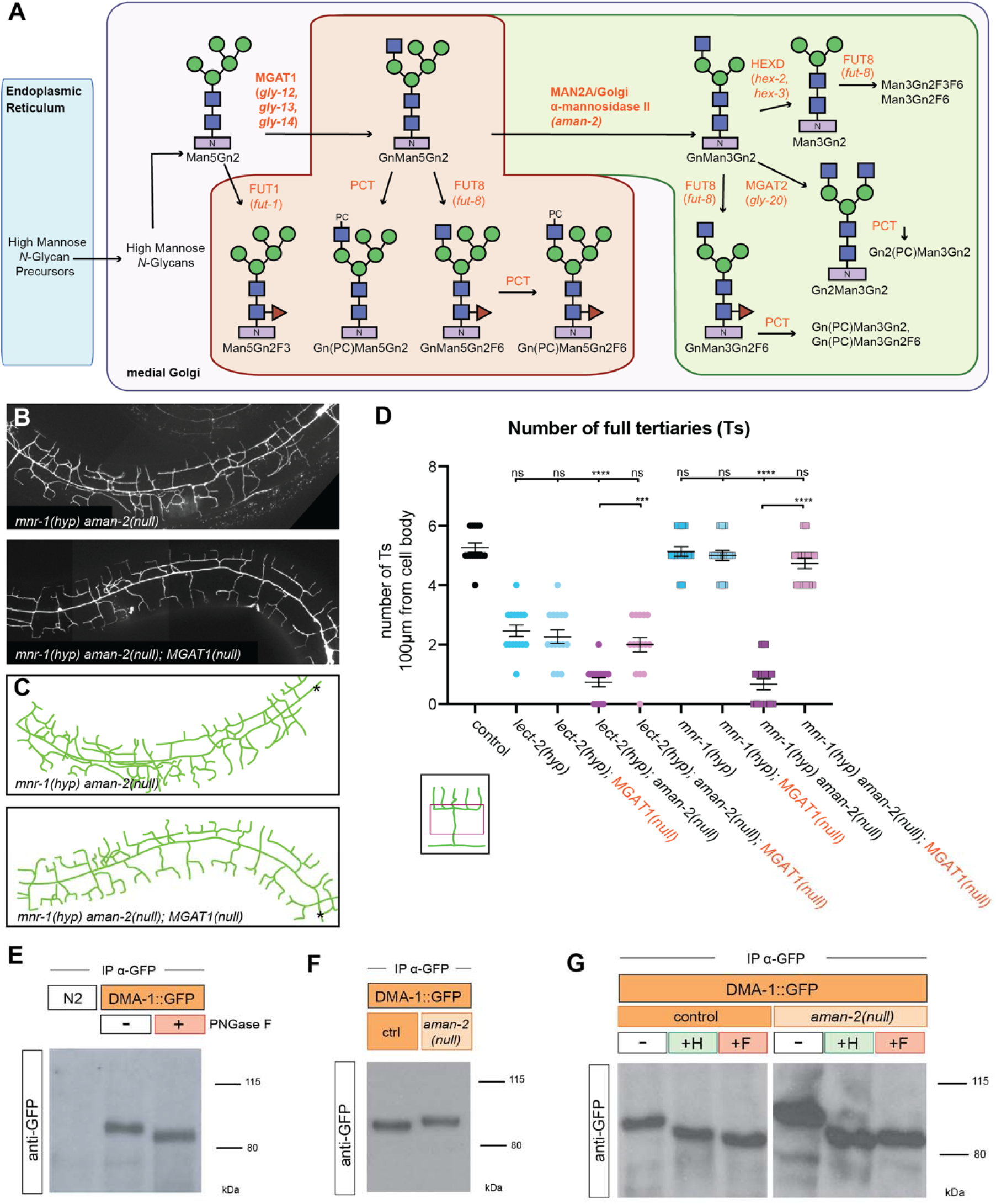
The presence of abnormal *N*-glycans in mutants of AMAN-2/Golgi alpha-mannosidase II results in defects in PVD arborization. (A) Schematic of the conserved *N*-glycosylation pathway in *C. elegans*. The blue box represents the Endoplasmic Reticulum, while the purple box represents the medial Golgi. Glycan residues are consistent with Figure 2A, with the addition of red triangles denoting fucose residues. Arrows and orange text represent enzymes. The green area marks wild type *N*-glycan chains whereas the red area represents abnormal *N*-glycan chains that arise in the absence of AMAN-2. The glycans in the green area are not formed in the absence of AMAN-2. (Man= mannose, Gn=N-acetylglucosamine, F=fucose, PC=phosphorylcholine, MGAT1=*N*-acetylglucosaminyltransferase I, FUT=fucosyltransferase, HEXD=hexosaminidase). (B-C) Fluorescent images (top) and tracings (bottom) of *mnr-1(dz213*) in an *aman-2*(*gk248486*) and an *aman-2*(*gk248486*); *MGAT-1*(*null*) background. An MGAT null mutant lacks the three *C. elegans* paralogs: *gly-12*, *gly-13*, and *gly-14*. PVD is visualized by the *wyIs581* transgene. The cell body is denoted with an asterisk. Anterior is to the left and dorsal is up in all panels. (D) Quantification of “Ts” of denoted genotypes. Data are represented as mean ± SEM. Statistical comparisons were performed using the Kruskal-Wallis test. Statistical significance is indicated (***p ≤ 0.001, ****p ≤ 0.0001, ns=not significant). n=15. (E) Western blot against GFP in *C. elegans* lysate expressing no transgenes (N2) and expressing DMA-1::GFP (*qyIs369*), after precipitating with anti-GFP antibody. The red boxed plus sign indicates that the lysate is treated with the PNGase F glycosidase. The downwards size shift reveals that *N*-glycan structures are present on DMA-1. Ladder is marked in kilodaltons (kDa). The GFP tag contains no *N*-glycosylation sites. (F) Western blot against GFP in *C. elegans* lysate DMA-1::GFP (*qyIs369*), after precipitating with anti-GFP antibody. Control indicates an otherwise wild type background as opposed to an *aman-2*(*gk248486*) null background. The downward size shift in the mutant reveals that loss of *aman-2* alters the identity of *N*-glycan structures on DMA-1. (G) Western blot against GFP in *C. elegans* lysate DMA-1::GFP (*qyIs369*), after precipitating with anti-GFP antibody. Control indicates an otherwise wild type background as opposed to an *aman-2*(*gk248486*) null background. The red boxed +F indicates that the lysate is treated with the PNGase F glycosidase, while the green boxed +H corresponds to the Endo H glycosidase, which cleaves high-mannose and hybrid type *N*-glycans. For complementary experiments using the Endo D glycosidase, which cleaves paucimannose type *N*-glycans, see Fig.EV4B. Size shifts indicate that some hybrid/high-mannose structures are present on DMA-1 (left), and that the *aman-2* mutant results in only hybrid/high-mannose structures on DMA-1 (right).

Since AMAN-2/Golgi alpha-mannosidase II canonically functions as an enzyme (Moremen, 2002; Shah *et al*, 2008), we next asked whether catalytic activity is required for its role in PVD dendrite branching. We approached this both genetically and pharmacologically. Prior studies showed that two highly-conserved aspartates are part of the conserved catalytic site in AMAN-2/Golgi alpha-mannosidases II (D306 and D443) and act sequentially to cleave off two mannose residues (Fig.2D) (Shah *et al*., 2008). We found that an AMAN-2 cDNA with both aspartates mutated, and hence likely catalytically dead, failed to rescue the defects in *lect-2*(*gk864764*); *aman-2*(*gk248486*) and *mnr-1*(*dz213*) *aman-2*(*gk248486*) double mutants (Fig. 2C). To address the possibility that mutating the catalytic residues compromised the stability or structure of AMAN-2, we took advantage of swainsonine, a compound that specifically inhibits Golgi alpha-mannosidase II (Lu *et al*, 2014). We found that exposing animals to swainsonine resulted in PVD defects that were indistinguishable from the effects of a null mutation in *aman-2*, either in combination with a partial loss of function allele of *lect-2*, or in wild type animals (Fig. 2E-G). In other words, the pharmacological inhibition of AMAN-2/Golgi alpha-mannosidase II activity resulted in the same phenotypic consequences as genetically inactivating or removing the enzyme. Collectively, these findings lead us to conclude that the catalytic activity of AMAN-2/Golgi alpha-mannosidase II is essential to support dendrite patterning in PVD. This further implies that *N*-glycosylation of a molecule expressed in PVD is crucial for normal dendrite arborization.

### The presence of abnormal *N*-glycans in *aman-2/Golgi alpha-mannosidase II* mutants results in defective PVD arborization

In eukaryotes, *N*-glycosylation is initiated in the Endoplasmic Reticulum (ER) with the synthesis of a 14-saccharide glycan on the phosphorylated polyisoprenol lipid dolichol-P-P (Stanley *et al*., 2015). Subsequently, the saccharide is transferred by a multiprotein complex termed oligosaccharyltransferase (OST) from dolichol-P-P to the aspartate within a NXS/T motif in nascent proteins as they are translocated into the ER (Stanley *et al*., 2015). As *N*-glycosylated proteins transit the Golgi, the glycans undergo a series of enzymatic modifications that add and remove specific sugar residues to lead to a wide array of possible *N*-glycan structures (Fig.3A) (Stanley *et al*., 2015). For example, in one of the earlier steps, the enzyme MGAT1 adds a *N*-acetylglucosamine residue to Man5Gn2 to form GnMan5Gn2 (Fig.3A). Genetically removing MGAT1 in mice results in complete loss of complex and hybrid *N*-glycans and early embryonic death, demonstrating that these *N*-glycans are essential for mammalian development (Ioffe & Stanley, 1994; Metzler *et al*, 1994). GnMan5Gn2 is the substrate for AMAN-2/MAN2A, which sequentially removes two mannose residues to form GnMan3Gn2 (Fig.3A) (Stanley *et al*., 2015). These reactions are followed by either removal or addition of additional sugars, or modification by a host of other conserved enzymes that lead to either complex or paucimannose *N*-glycans (Fig.3A). To determine which specific *N*-glycans are missing in *aman-2* mutants, and are therefore required for branching of PVD dendrites, we systematically tested whether mutations in any of the genes downstream of *aman-2* (including *hex-2/hexosaminidase*, *hex-3/hexosaminidase*, *fut-8/FUT8 Fucosyltransferase*, and *gly-20/MGAT II*) would also enhance the partial *lect-2* loss of function allele. We found that removing the genes encoding these enzymes alone, or in combination, did not enhance the partial *lect-2* loss of function allele (Fig.EV3A). Mutating MGAT1/*N*-acetylglucosaminyltransferase-I (in worms encoded by three paralogous genes *gly-12, gly-13, gly-14)* (Chen *et al*, 1999; Chen *et al*, 2003), which acts immediately before AMAN-2/Golgi alpha-mannosidase II, also showed no effects (Fig.EV3B). Collectively, these findings suggest that no lack of specific *N*-glycans downstream of MGAT1, or of AMAN-2, alone are responsible for the observed defects in PVD dendrites.

Previous structural studies of *N*-glycans in *aman-2*(*tm1078*) null mutant animals (Paschinger *et al*, 2006) established that loss of *aman-2/Golgi alpha-mannosidase II* in *C. elegans* caused (1) a loss of the normal products of AMAN-2, including complex *N*-glycans (Fig.3A, shaded in green) and (2) a buildup of GnMan5Gn2, the substrate of AMAN-2 (Fig.3A)(Paschinger *et al*., 2006). This GnMan5Gn2 intermediate was found to serve as substrate for enzymes further downstream (including *FUT-8/Fut8 Fucosyltransferase* and *PCT/*Phosphorylcholine-transferase) leading to the appearance of abnormal *N*-glycans, not normally present in wildtype animals (Fig.3a, shaded in red) (Paschinger *et al*., 2006). To determine whether the defects in PVD branching were caused by the absence of wild type *N*-glycans, or the presence of abnormal GnMan5Gn2 *N*-glycans, we mutated both MGAT1 and AMAN-2 in the partial loss of function backgrounds of *lect-2* and *mnr-1*. The prediction was that, if the enhancement of the partial loss of function alleles *lect-2* or *mnr-1* by loss of *aman-2* is caused by abnormal *N*-glycans, then removal of MGAT1, the preceding enzyme would suppress that enhancement. Indeed, loss of MGAT1 did suppress the enhanced phenotypes in *lect-2*; *aman-2* and *mnr-1*; *aman-2* double mutants (Fig.3B-D). These data indicate that one or more structurally abnormal *N*-glycans with a terminal *N*-acetylglucosamine are responsible for the observed defects in PVD patterning.

### DMA-1/LRR-TM *N*-glycosylation is changed in *aman-2/Golgi alpha-mannosidase II* mutants

Since we demonstrated that (1) *aman-2/Golgi alpha-mannosidase II* genetically interacts with the Menorin pathway, and (2) AMAN-2/Golgi alpha-mannosidase II activity is required cell-autonomously in PVD to regulate branching, we hypothesized that AMAN-2/Golgi alpha-mannosidase II directly regulates *N*-glycans on at least one component of the Menorin complex in PVD. Treatment of whole worm lysates with the bacterial PNGase F glycosidase, which cleaves all *N*-glycans from Asn, resulted in distinct downward shifts in molecular weight of both DMA-1 and KPC-1, indicating that *N*-glycans were present on both proteins *in vivo* and had been removed (Fig.3E,EV4C). Thus, both PVD-expressed proteins, DMA/LRR-TM and KPC-1/Furin, are *N*-glycosylated, consistent with a previous report for DMA-1/LRR-TM (Feng *et al*, 2020). Another cell-autonomous factor and possible candidate, HPO-30/Claudin (Smith *et al*, 2013), contains no predicted *N*-glycan consensus motifs.

We next determined whether loss of AMAN-2 resulted in altered *N*-glycosylation. Interestingly, the absence of *aman-2/Golgi alpha-mannosidase II* resulted in a clear increase in DMA/LRR-TM molecular weight, whereas the size of KPC-1/Furin remained unaffected (Fig.3F,EV4C). The upward shift in DMA-1 size following the loss of *aman-2* is consistent with our genetic data establishing that the presence of larger, abnormal GnMan5Gn2 *N*-glycans gives rise to the PVD mutant phenotype. To determine what types of *N*-glycans are attached to DMA-1 in wild type and *aman-2* mutant backgrounds, we treated lysates with additional endoglycosidases: Endo H, which cleaves hybrid/high-mannose *N*-glycans, and Endo D, which cleaves only paucimannose *N*-glycans. Based on the size of the shifts observed, we conclude that in wildtype animals, DMA-1 possesses primarily, but not exclusively, hybrid/high-mannose *N*-glycans with a smaller amount of paucimannose *N*-glycans. In contrast, in the absence of AMAN-2, all *N*-glycans on DMA-1/LRR-TM were converted to hybrid/high-mannose type (likely GnMan5Gn2-derived *N*-glycans) with no or little detectable paucimannose *N*-glycans (Fig.3G,EV4A,B). Additionally, proteomic studies identified *C. elegans* SAX-7/L1CAM and LECT-2/Chondromodulin II as glycoproteins (Kaji *et al*, 2007), which we confirmed by a shift in molecular weight upon digestion of all *N*-glycans by PNGase F (Fig.EV4D,E). However, in the absence of *aman-2/Golgi alpha-mannosidase II*, neither SAX-7/L1CAM and LECT-2/Chondromodulin II displayed obvious changes in molecular weight, suggesting that they did not carry abnormal *N*-glycans in an *aman-2* mutant background. Collectively, our data show that among the *N*-glycosylated proteins of the Menorin complex, only DMA-1/LRR-TM was significantly affected by the loss of *aman-2/Golgi alpha-mannosidase II* and carried altered *N*-glycans in *aman-2* null animals.

### AMAN-2/Golgi alpha-mannosidase II modulates *N*-glycans on DMA-1/LRR-TM to regulate PVD morphogenesis

The DMA-1/LRR-TM receptor contains four predicted *N*-glycosylation motifs, all of which reside in leucine rich repeats (Fig.4A). To establish whether the *N*-glycosylation of DMA-1/LRR-TM is essential for its role in PVD dendrite branching, we mutated all four sites, alone and in combinations. Using CRISPR/Cas9-based genome editing, we converted the asparagine residues of the four predicted *N*-glycan attachment sites to glutamine to maintain chemical similarity but eliminate the possibility of *N*-glycosylation. We found that only when abolishing predicted *N*-glycan attachment site 4 (N386), alone or in combination with other sites, was PVD quaternary branching compromised (Fig.4B). These results reveal that *N*-glycosylation of DMA-1/LRR-TM is required for its role in PVD patterning of quaternary branches and highlight the importance of N386 within a membrane proximal LRR repeat. We then assessed whether abolishing *N*-glycan attachment sites in an *aman-2/Golgi alpha-mannosidase II* loss of function background had any effects on dendrite patterning. Analysis of different site-specific mutations revealed that some *N*-glycosylation sites on DMA-1/LLR-TM enhance the severity of PVD dendrite branching defects in an *aman-2* mutant background (Fig.4C-D). The results in an *aman-2* mutant background suggest that having some type of *N*-glycan on site 4 of DMA-1/LRR-TM, even if abnormal, is better than having no *N*-glycan at all (cf. S3 and S123); second having such abnormal *N*-glycans on sites 1-3 further compromises DMA-1 function during PVD development (cf. S4 in control vs *aman-2* mutant background, Fig.4B,C). Lastly, the mutant phenotype resulting from a presumptive loss of all *N*-glycans on DMA-1 (S1234) is enhanced in an *aman-2* mutant background, suggesting that *N*-glycosylated proteins other than DMA-1 may serve additional functions during PVD morphogenesis or that abnormal *N*-glycans on cryptic *N*-glycosylation sites further compromise function (Fig.4C).

**Figure 4.**
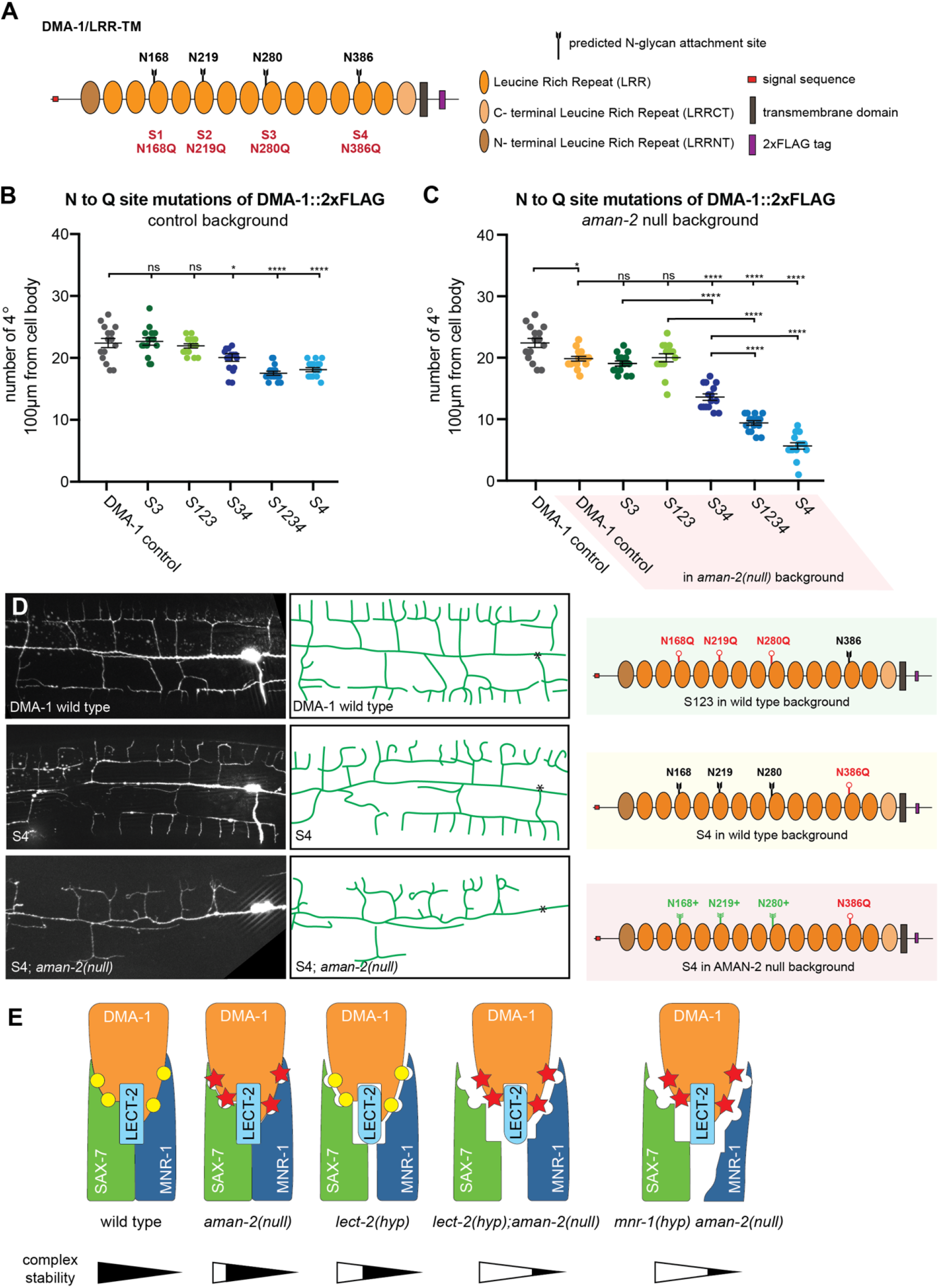
AMAN-2/Golgi alpha-mannosidase II modulates *N*-glycans on DMA-1/LRR-TM to regulate PVD morphogenesis. (A) Schematic of DMA-1/LRR-TM with mutated *N*-glycan attachment sites S1-S4 indicated. (B) Quantification of the number of quaternary branches in DMA-1::2xFLAG control animals and animals with combinations of DMA-1::2xFLAG *N*-glycan attachment sites mutated. Note that DMA-1::2xFLAG control animals display a slightly reduced number of quaternary dendrites compared to wild type animals (Fig.EV3C). Data are represented as mean ± SEM. Statistical significance was calculated using the Kruskal-Wallis test and is indicated (****p ≤ 0.0001, ns=not significant). n = 15 for all genotypes. (C) Quantification of the number of quaternary branches in DMA-1::2xFLAG control animals alone and in combination with an *aman-2*(*gk248486*) null mutant (shaded in red). Control data is identical as in (B) and shown for comparison only. Data are represented as mean ± SEM. Statistical significance was calculated using the Kruskal-Wallis test and is indicated (*p ≤ 0.05, ****p ≤ 0.0001, ns=not significant). n = 15 for all genotypes. (D) Fluorescent images (left) and tracings (center) of PVD in animals of denoted backgrounds (*aman-2(null*) is *gk248486*). S4 corresponds to the endogenous alteration of *N*-glycan attachment site N386 of DMA-1. PVD is visualized by the *dzIs117* transgene. The cell body is marked with an asterisk. The schematics on the right detail the molecular context of the indicated backgrounds. Green indicates a relatively normal PVD structure, yellow a slightly compromised structure, and red a heavily defective arborization. Black attachments indicate wild type, red indicate blocked sites, and green indicates larger abnormal *N*-glycans chains. (E) The proposed model of how the loss of *aman-2* can modulate *N*-glycans on DMA-1, and potentially the binding of the Menorin complex as a whole. Yellow circles represent normal *N*-glycan chains, while red stars represent abnormal *N*-glycans produced in an *aman-2* mutant background. Putative complex stability is indicated below.

## DISCUSSION

While prior studies established the importance of *N*-glycosylation during neuronal development, our studies establish an important role for specific classes of *N*-glycans in mediating neuronal development, and specifically dendrite patterning. They suggest that the DMA-1/LRR-TM receptor in PVD must be decorated with specific hybrid/high-mannose or paucimannose *N*-glycans as a result of AMAN-2 activity and that these specific *N*-glycans are important for functioning of the Menorin complex in PVD dendrite morphogenesis. A possible explanation is that the *N*-glycans on specific *N*-glycosylation sites in DMA-1/LRR-TM function permissively to maintain high affinity binding of DMA-1/LRR-TM to other members of the Menorin complex through specific *N*-glycans (Fig.4E). This interaction could be compromised by the formation of abnormal glycans in the absence of AMAN-2 leading to destabilization of the complex. Therefore, our genetic and biochemical data, together with analytical data of *N*-glycans in *aman-2/Golgi alpha-mannosidase II* mutants (Paschinger *et al*., 2006), underscore the importance of AMAN-2/Golgi alpha-mannosidase II as a linchpin for the creation of complex and paucimannose *N*-glycans, and avoidance of larger, abnormal hybrid and high-mannose-type *N*-glycans. Since metabolite availability can influence *N*-glycan synthesis and flux (reviewed in (Dennis *et al*, 2009)), these findings also raise the possibility that environmental factors can intersect with intrinsic genetic programs to regulate extracellular adhesion complexes during neural development by modulating *N*-glycosylation.

Previous studies demonstrated that *N*-glycosylation is important for folding and surface localization of cell adhesion molecules and axon guidance factors, such as L1CAMs and ephrins, respectively (Sekine *et al*., 2013; Medina-Cano *et al*., 2018; Mire *et al*., 2018). On the other hand, *in vitro* experiments suggested that *N*-glycans can regulate protein-protein interactions of cell adhesion molecules (Fogel *et al*, 2010; Labasque *et al*, 2014). Our findings demonstrate *in vivo*, that not only *N*-glycans *per se*, but that specific classes of *N*-glycan structures are important to modulate cell-cell signaling, and possibly, receptor-ligand binding and complex formation. This is reminiscent of the role of *O*-fucose glycans on the Notch receptor extracellular domain, which affect its signaling and ligand interactions (Moloney *et al*, 2000). Given that over 70% of proteins transiting the secretory pathway are *N*-glycosylated (Apweiler *et al*., 1999), these findings raise the possibility that specific *N*-glycan structures are important determinants to regulate the interactions of extracellular complexes during nervous system development more broadly and could contribute to the specificity that is required in the nervous system. In this context it is interesting to note that over 70 congenital disorders of glycosylation have been described that affect genes in the *N*-glycosylation pathway (Freeze, 2006; Ng & Freeze, 2018), of which many are associated with intellectual disability or other neurological symptoms (Jaeken & Peanne, 2017; Chang *et al*., 2018). While no mutations in Golgi alpha-mannosidase II in humans have been described to date, it is conceivable that such mutations exist, and may result in neurological phenotypes. Regardless, our studies provide the conceptual framework for studies into developmental defects of the nervous system in mutants of genes involved in *N*-glycosylation and this growing class of congenital disorders.

## ACKNOWLEDGEMENTS

We thank Yehuda Salzberg, Pamela Stanley, Robert Townley, Peri Kurshan and members of the Bülow laboratory for comments on the manuscript and helpful discussions during the course of this work. We thank Kang Shen, Iain Wilson, Shohei Mitani for reagents, and Yuji Kohara for the *yk11g705* cDNA clone. We are grateful to Meera Trivedi for sharing the *dzIs117* strain prior to publication. Some strains were provided by the Caenorhabditis Genome Center (funded by the NIH Office of Research Infrastructure Programs P40 OD010440). This work was supported by grants from the National Institute of Health (NIH): R01NS096672 and R21NS111145 to HEB; F31NS100370 to MR; T32GM007288 and F31HD066967 to CADB; P30HD071593 to Albert Einstein College of Medicine. NJRS was the recipient of a Colciencias-Fulbright Fellowship and HEB of an Irma T. Hirschl/Monique Weill-Caulier research fellowship.

## AUTHOR CONTRIBUTIONS

Conceptualization Ideas: MR, HEB; Validation: MR; Formal Analysis: MR, HEB; Investigation: MR; Resources: NJRS, CADB; Writing – Original Draft: MR; Writing – Review & Editing: MR, NJRS, CADB, HEB; Visualization Preparation: MR; Project Administration: HEB; Funding Acquisition: MR, NJRS, CADB, HEB.

## MATERIALS AND METHODS

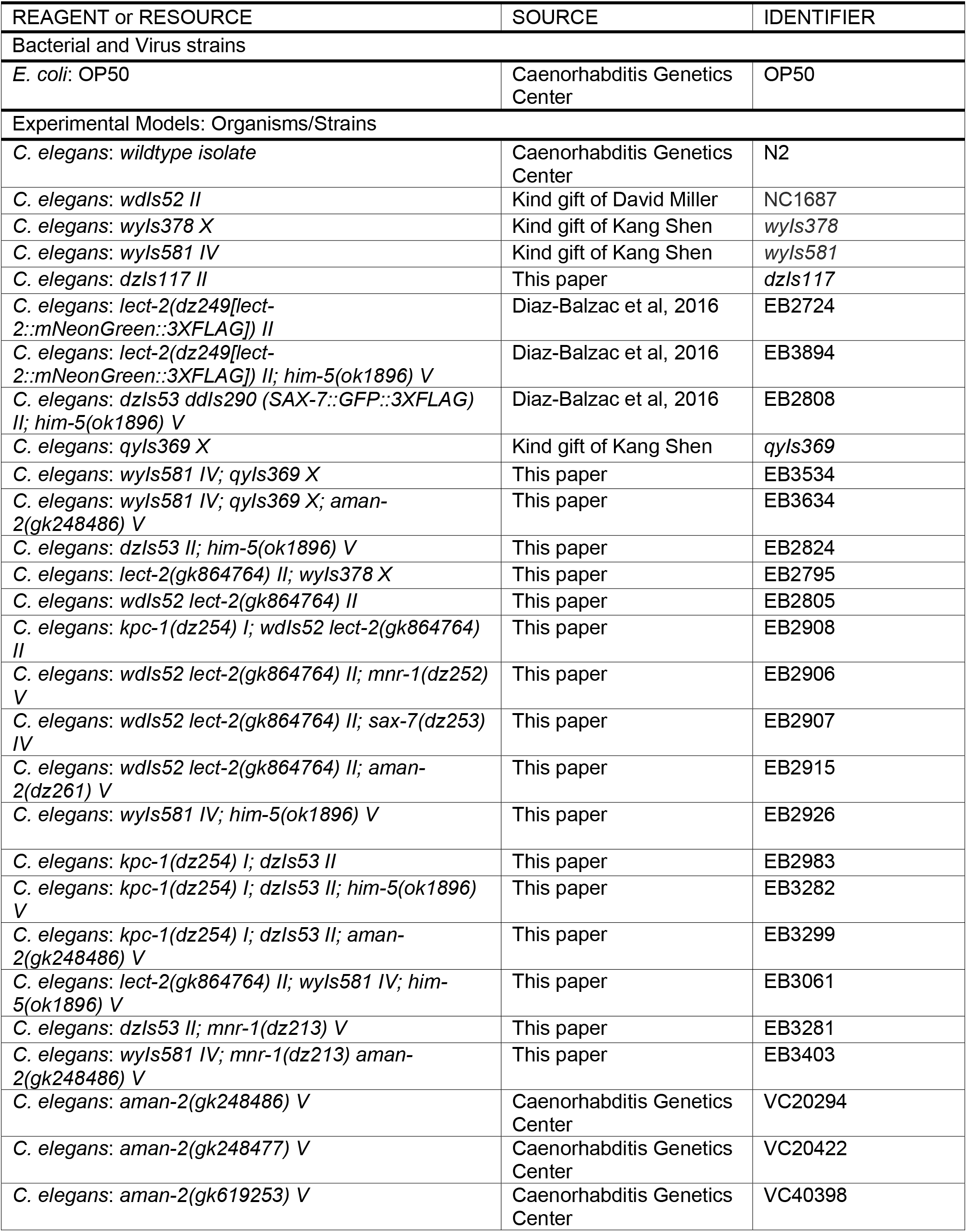
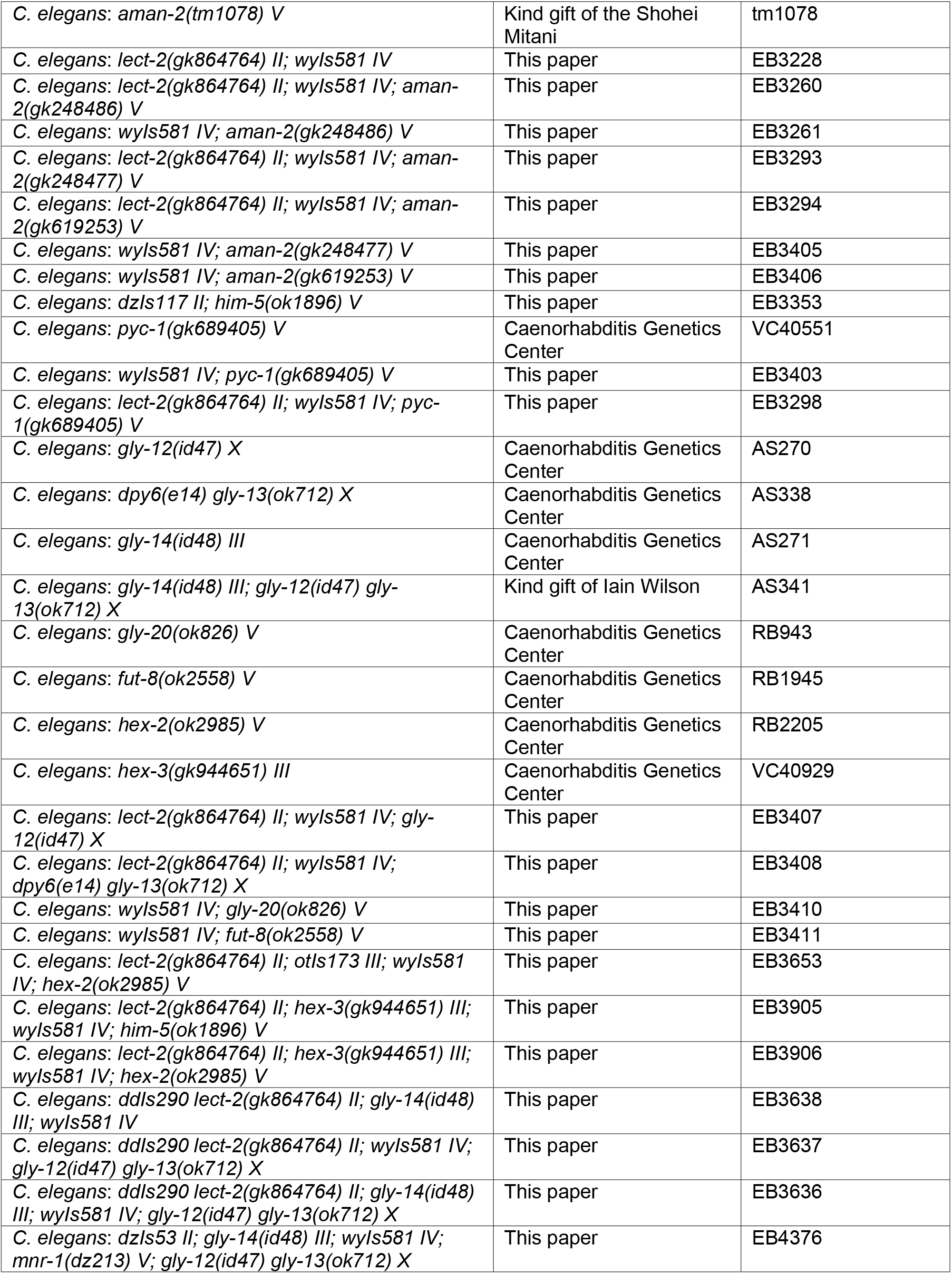
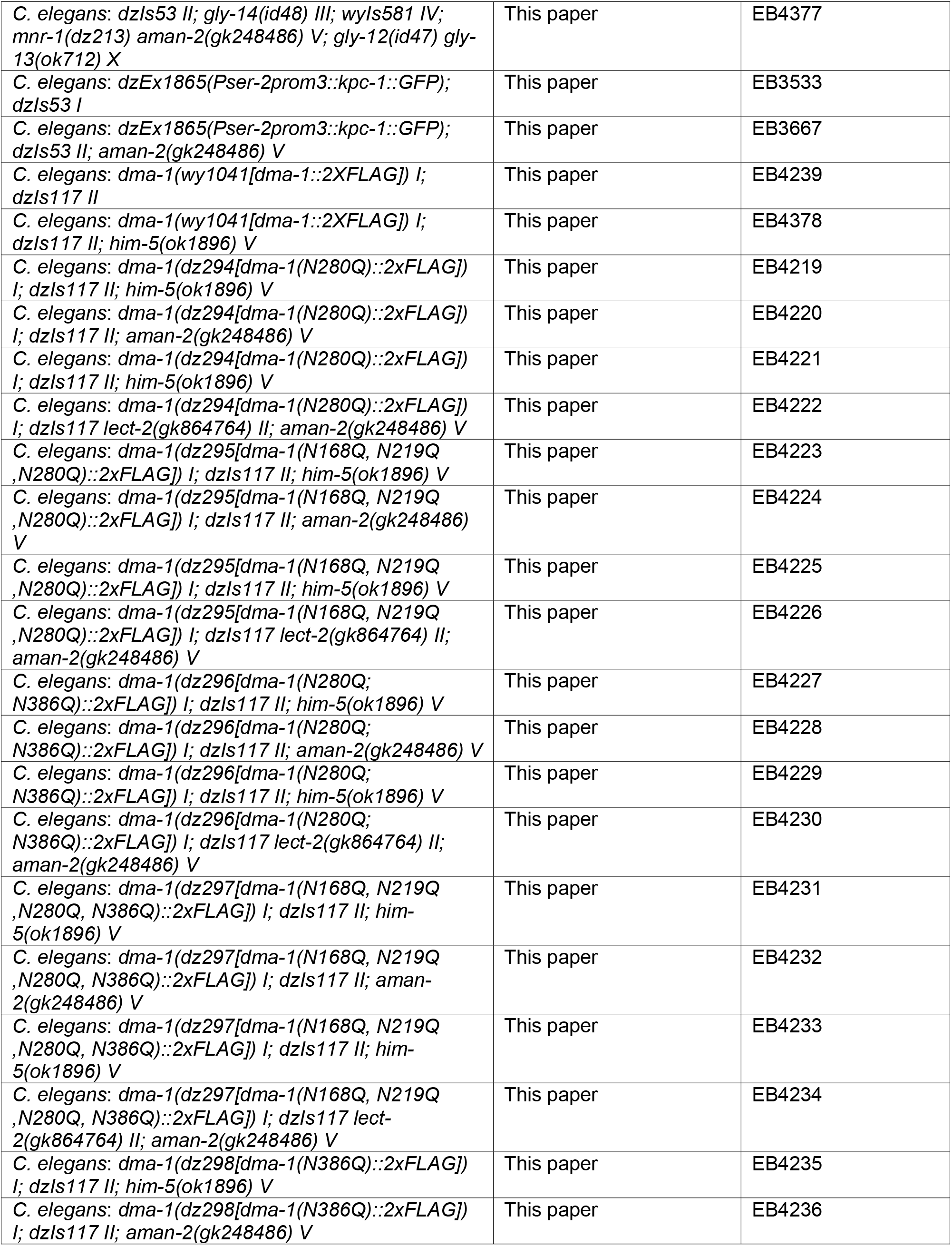
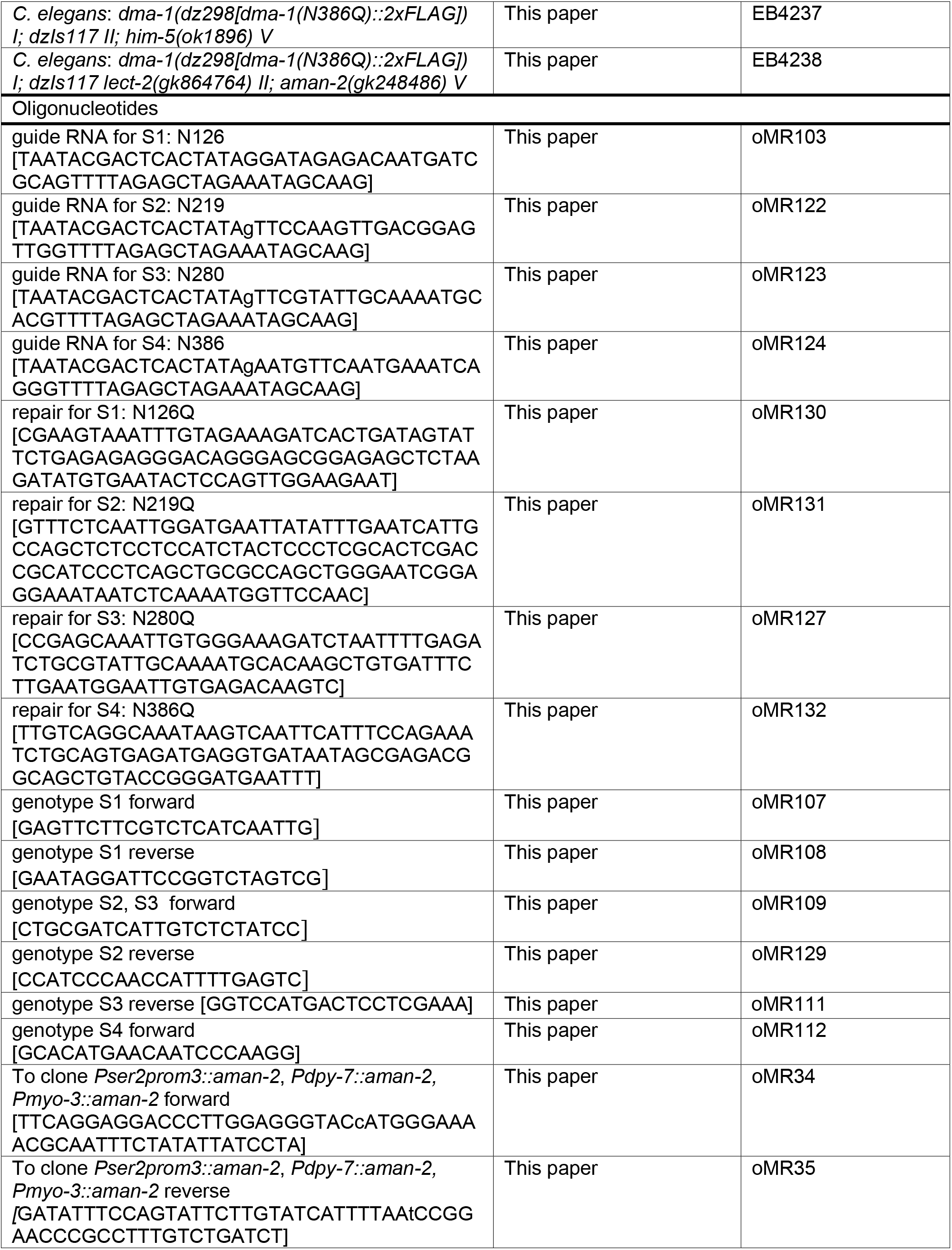
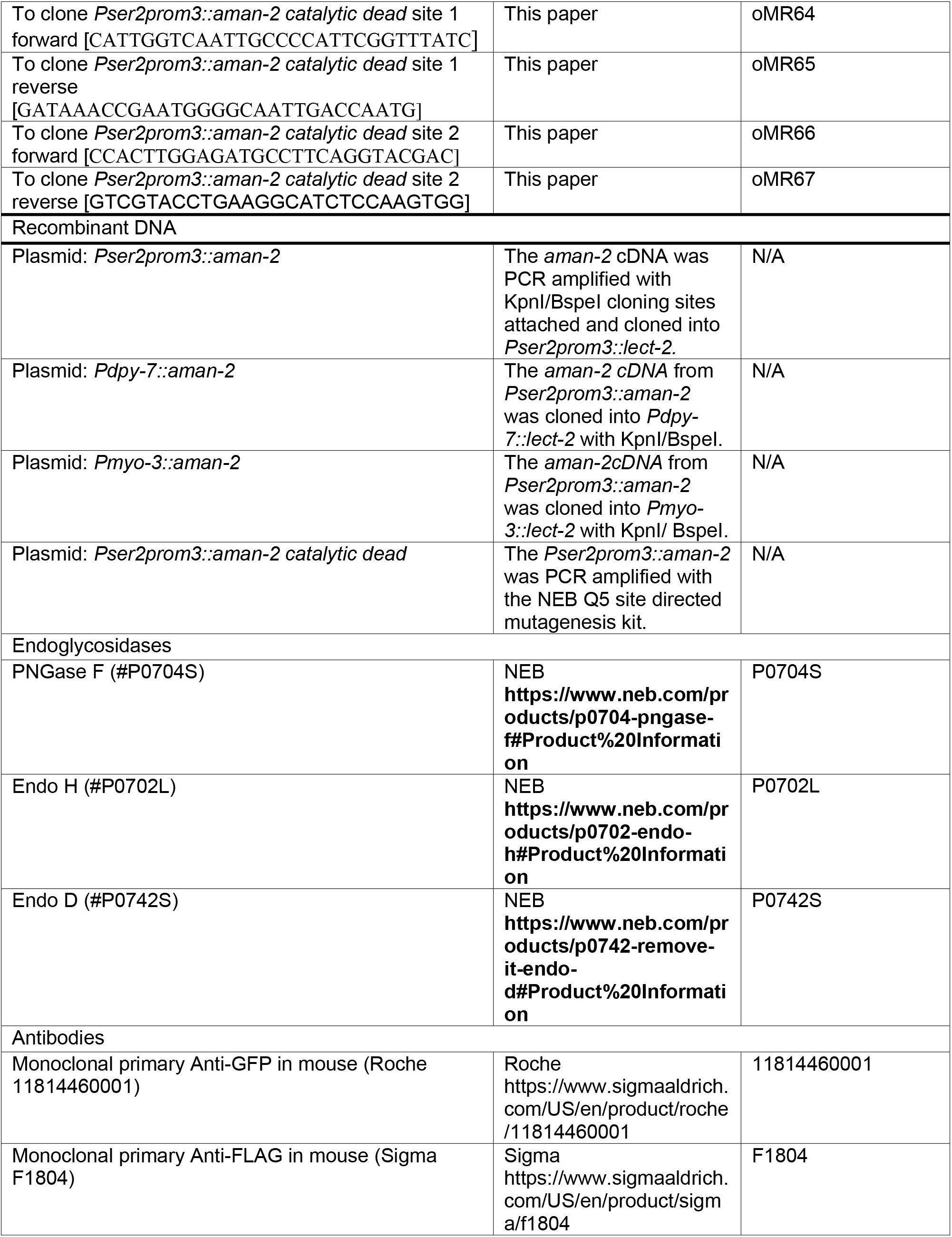
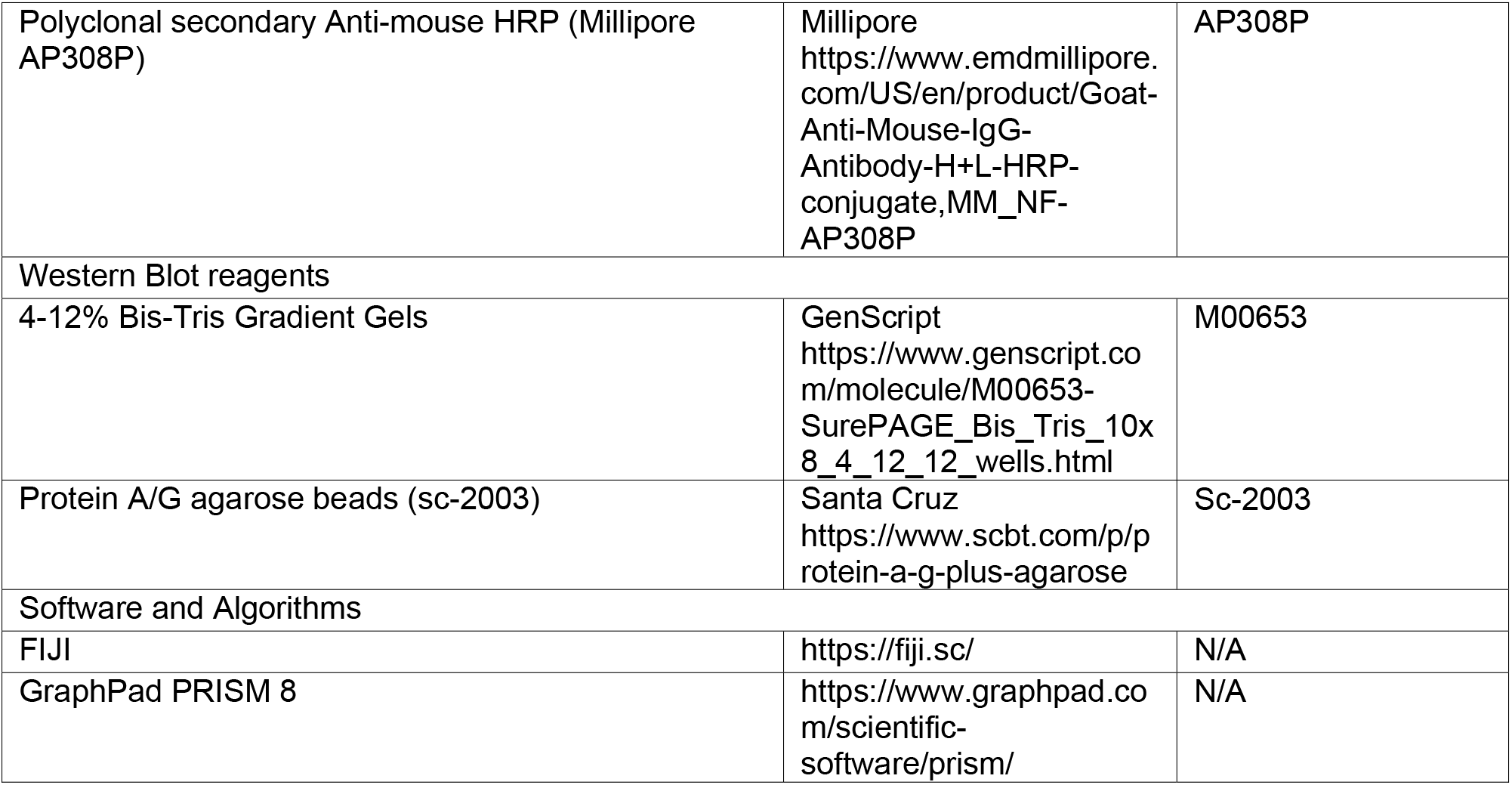

### *C. elegans* handling

All strains were maintained using standard methods (Brenner, 1974) and experiments were performed at 20°C, except where indicated otherwise. Phenotypic analysis was performed in 1-day-old adults, with no more than 4-5 eggs present. For details and a complete list of strains used and generated in this study, see resources table.

### Cloning of mutant alleles

The *dz261* allele was obtained from a forward genetic screen for modifiers of the *lect-2*(*gk864764*) hypomorphic allele. Using a combination of whole genome sequencing and single nucleotide polymorphism mapping (Minevich *et al*, 2012), we narrowed down the region to a 5Mb interval (11MB-16Mb on chromosome V (Fig.EV1A). This region contained 7 polymorphisms with predicted functional consequences. We injected seven fosmids in pools and found that only the pool which contained a fosmid covering *aman-2* resulted in rescue (Fig.EV1B). In addition, we obtained three nonsense alleles in *aman-2*(*gk248486*, *gk248477*, *gk619253*) from the Million Mutation Project (Thompson *et al*, 2013) and one deletion allele (*tm1078*, kind gift from the Mitani lab).

### Details of genetic screen and cloning

The *lect-2*(*gk864764*) hypomorphic strain was treated with EMS in accordance with standard chemical mutagenesis protocols (Kutscher & Shaham, 2014) and F1 progeny were scored for enhancement, suppression, or modification of PVD branching phenotype. A SNP-mapping-WGS (Doitsidou *et al*, 2016) approach was used to map *aman-2*(*dz261*) between 10Mb and 16Mb of chromosome V. This region contained seven candidates with nonsense, missense, frameshift or splice site mutations, including one in *aman-2*. The *dz261* mutation was further confirmed by Sanger sequencing of the original isolate, identifying a nonsense mutation W237Opal in the *aman-2* locus.

### Heterologous rescue of PVD dendrite branching defect

The aman-2 cDNA was cloned under control of heterologous promoters: hypodermal *Pdpy-7* (Gilleard *et al*, 1997), body wall muscle *Pmyo-3* (Okkema *et al*, 1993), and PVD *Pser2prom3* (Tsalik *et al*, 2003). All constructs were injected at 5 ng/μl into *wdIs52 II; him-5(ok1896) V* together with the *Pmyo-2::mCherry* marker at 50 ng/μl and *pBluescript* at 50 ng/μl. Males from transgenic lines were then crossed into *lect-2*(*gk864764*) *II*; *wyIs581 IV*; *aman-2*(*gk248477)V* and *wyIs581 IV*; *mnr-1*(*dz213*) *aman-2*(*gk248486*) *V*.

### Cloning constructs and transgenesis

To assemble tissue specific expression constructs used for heterologous rescue experiments, the *aman-2* cDNA clone *yk11g705* (kind gift of Yoji Kohara) and cloned under control of the following promoters: PVD *ser2prom-3* (Tsalik *et al*., 2003), hypodermal *dpy-7p* (Gilleard *et al*., 1997), body wall muscle *myo-3p* (Okkema *et al*., 1993). These constructs were injected at 5 ng/μl together with *myo-2p::mCherry* as an injection marker at 50 ng/μl, and *BlueScript* as DNA filler. Point mutants in the *aman-2* cDNA were introduced by site-specific mutagenesis (NEB Q5 Site-Directed

Mutagenesis). All plasmids contained the *unc-54* 3’UTR.

### Pharmacology

Experiments in which the activity of AMAN-2 was blocked pharmacologically were performed with the compound swainsonine (1 mg swainsonine #16860 vials, Fisher Scientific catalog #NC1670046). Dosage experiments were performed from 50-500μm of swainsonine in dissolved in agar of NGM plates, with 300μm being sufficient to elicit phenotypes in PVD. After drying for 24 hours, 200μL of OP50 *E. coli* was seeded onto each plate, and 5 young adult worms were left to self-fertilize and lay eggs. The F1 generations were analyzed, imaged, and quantified. DMSO was used a control in plates not treated with swainsonine.

### Molecular Biology

Immunoprecipitation and Western blot analyses of glycoproteins were performed using standard SDS-PAGE methods. Whole *C. elegans* lysates were prepared in RIPA buffer and sonicated in a Biorupter benchtop waterbath sonicator for 15 minutes. Lysates were treated with 1 unit at temperatures as indicated in NEB protocols (linked in Key Resources Table) endoglycosidases PNGase F, Endo H, or Endo D where specified. Overnight immunoprecipitation of lysates with anti-GFP antibody prior to SDS-PAGE was performed for proteins with low expression levels (DMA-1::GFP and KPC-1::GFP).

### Immunoprecipitation and Western blot Analysis

Five full plates of DMA-1::GFP and KPC-1::GFP tagged worms were washed in RIPA buffer pH7.0 and lysed for 15 minutes in a Biorupter water bath used previously published methods of whole worm protein extraction (Li & Zinovyeva, 2020). Post lysis, 20 uL of Protein A/G Plus Agarose beads (Santa Cruz sc-2003) and 1 uL of anti-GFP antibody (Roche 11814460001) were used to pull down DMA-1::GFP and KPC-1::GFP overnight at 4C. Ten gravid adult SAX-7::GFP::FLAG animals and twenty LECT-2::mNG::FLAG animals were sufficient to see robust expression post Western Blot. These samples were boiled and loaded directly into the gels. Gradient gels (4-12% GenScript) were used in all experiments. For all anti-FLAG blots, a concentration of 1:800 anti-Flag (Sigma F1804) and 1:5000 anti-mouse HRP (Millipore AP308P) were used. For all anti-GFP blots, a concentration of 1:500 anti-GFP (Roche 11814460001) and 1:5000 anti-mouse HRP (Millipore AP308P) were used.

#### CRISPR/Cas9 mediated gene editing

CRISPR-Cas9 constructs were designed and dpy-10 co-crispr protocol followed as previously described (Dickinson & Goldstein, 2016) to make single point mutations of predicted *N*-glycan attachment Asparagine residues. Predicted sites were identified using NetNGlyc 1.0 Server (Gupta & Brunak, 2002). A battery of guideRNAs were designed to direct Cas9 cuts near sites of interest, and homologous repair template oligomers were designed to mutate Asparagine residues to Glutamine. All plasmids were delivered via microinjection in the gonads of animals. Strains with combinations of mutated sites were generated by sequential injections and/or multiple simultaneous edits. Note, that all edits were made in strains with the C-terminus of DMA-1 already tagged with a 2XFLAG immunotag before the PDZ binding domain (Dong *et al*., 2016). Strains EB4219 through EB4238 in the Key Resources Table were obtained using these methods.

### Imaging

Fluorescent images were captured in live *C. elegans* using a Plan-Apochromat 40×/1.4 or 63x/1.4 objective on a Zeiss Axioimager Z1 Apotome. Worms were immobilized using 1 mM Levamisole and *Z* stacks were collected. Maximum intensity projections were used for further analysis and tracing of dendrites. For quantification of branching, 1-day-old adults were mounted onto slides and immobilized with 1mM Levamisole. In both cases of capturing images and counting, and counting live on the microscope, the number of “Ts” (secondary and tertiary branches), “Os,” (self-avoidance defects), and/or quaternary branches within 100 μm of the primary branch anterior to the cell body were quantified.

### Quantification and statistical analysis

Statistical comparisons were conducted on Prism 8 GraphPad Software using Mann-Whitney, Kruskal-Wallis, Z-, or two-sided ANOVA tests as appropriate. Statistical significance is indicated as ns, not significant; *p≤0.05; **p≤0.01; ***p≤0.001 and ****p≤0.0001. This study includes no data deposited in external repositories.

## EXPANDED VIEW FIGURES

**Figure EV1.**
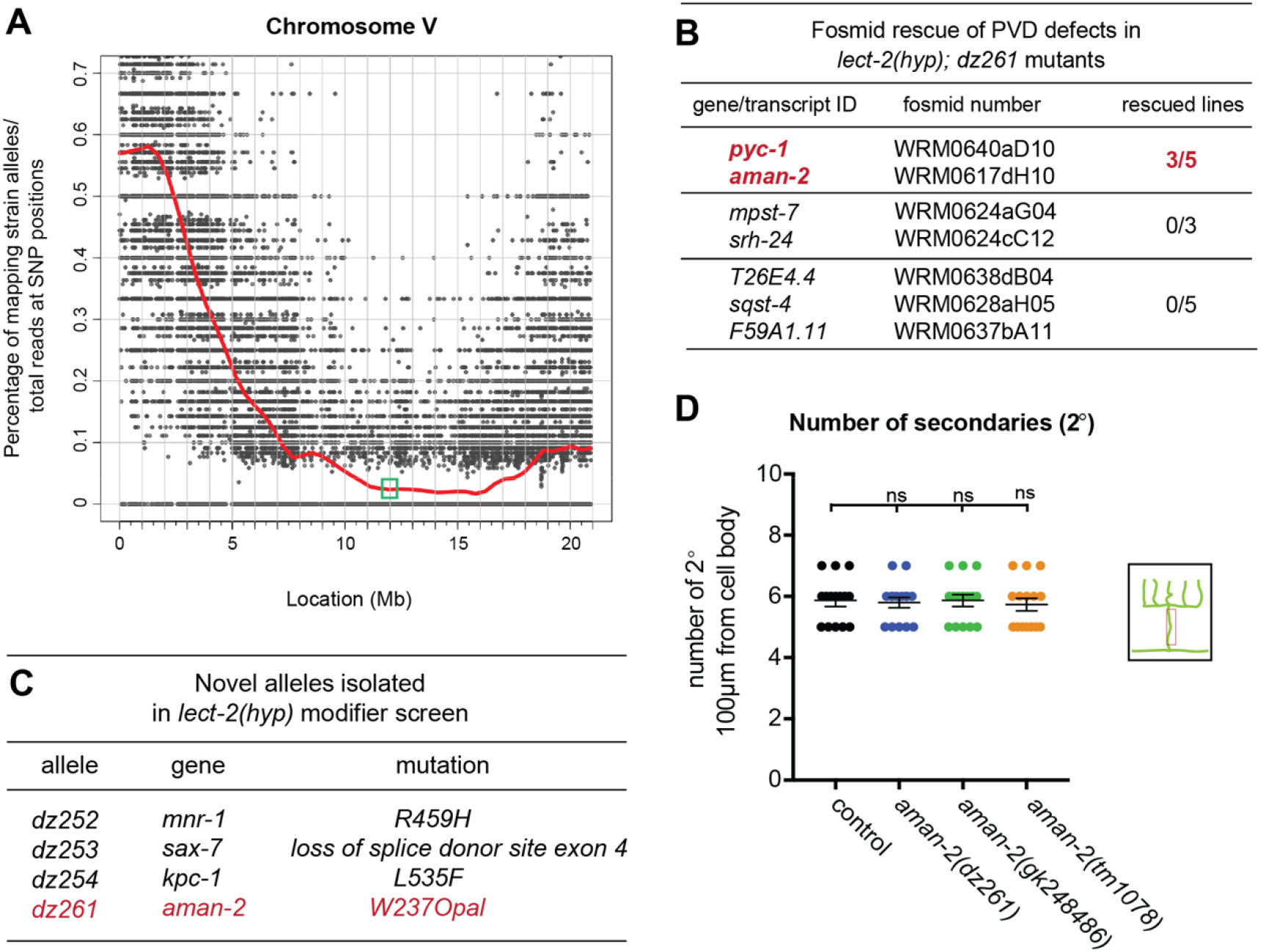
Results of modifier screen and identification of AMAN-2/Golgi Alpha-mannosidase II. (A) Graph of single nucleotide polymorphism (SNP) results after whole genome sequencing of the *dz261* allele (Doitsidou *et al*., 2016). Axes are denoted above. The green box shows the genomic position of *aman-2* (11.92-11.93 Mb on Chromosome V). (B) Table showing the candidate genes selected and tested as a result of the SNP data in (A). Pools of fosmids covering the regions of indicated transcripts were injected into the *lect-2(hyp) dz261* double mutant. Only the pool containing a fosmid including *aman-2* showed rescue. *pyc-1* was eliminated because a null allele (*gk689405*) failed to enhance the *lect-2(hyp*). (C) Table summarizing different additional alleles isolated in the *lect-2(gk864764*) hypomorph modifier screen and their molecular lesions. (D) Quantification of the number of secondary branches (indicated in schematic) in various *aman-2* mutant alleles and in wild type control animals. Data are represented as mean ± SEM. Statistical significance was calculated using the Kruskal-Wallis test and is indicated (ns= no significance). n = 15 for all genotypes.

**Figure EV2.**
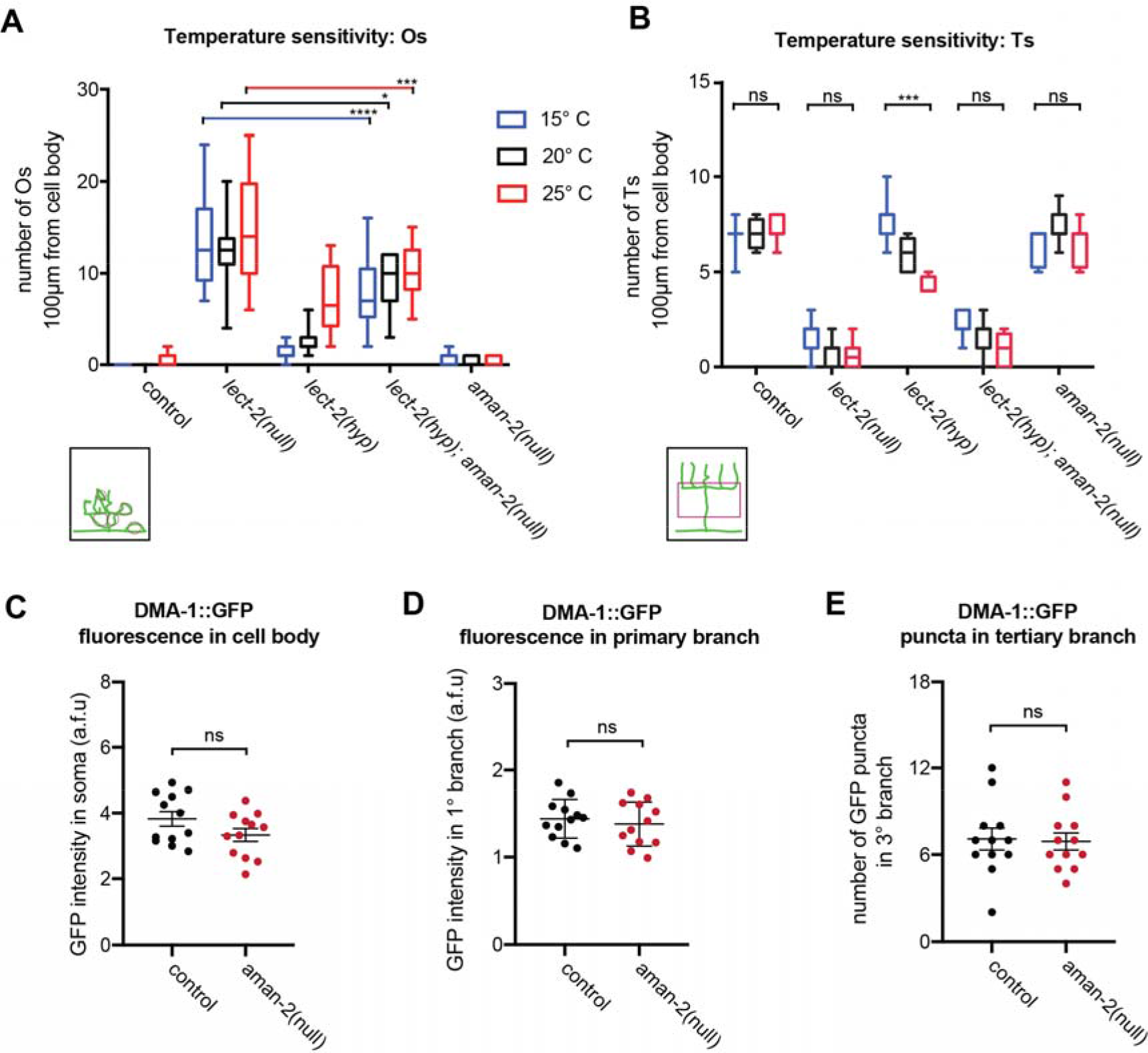
The requirement of AMAN-2 in PVD may be specific rather than global. (A) Quantification of “Os” (overlapping branches as shown in schematic) of denoted genotypes at different temperatures. Data are represented as mean ± SEM. Statistical comparisons were performed using two-way ANOVA tests. Statistical significance is indicated (*p ≤ 0.05, ***p ≤ 0.001, ****p ≤ 0.0001, ns=not significant). n > 11 for all groups. (B) Quantification of “Ts” of denoted genotypes at different temperatures. Data are represented as mean ± SEM. Statistical comparisons were performed using two-way ANOVA tests. Statistical significance is indicated (***p ≤ 0.001, ns=not significant). n > 11 for all groups. (C) Quantification of DMA-1::GFP fluorescence in control and *aman-2*(*gk248486*) animals. GFP intensity is quantified in arbitrary fluorescent units by dividing the fluorescent area of the soma by the background. Data are represented as mean ± SEM. Statistical comparisons were performed using the Mann-Whitney test. (ns=not significant). n=12. (D) Quantification of DMA-1::GFP fluorescence in control and *aman-2*(*gk248486*) animals. GFP intensity is quantified in arbitrary fluorescent units by measuring the fluorescence along the primary dendrite, up to 60μm anterior from the cell body. Data are represented as mean ± SEM. Statistical comparisons were performed using the Mann-Whitney test. (ns=not significant). n=12. (E) Quantification of DMA-1::GFP fluorescent puncta in control and *aman-2*(*gk248486*) animals. Puncta in the tertiary branches 60μm anterior to the cell body were counted. Data are represented as mean ± SEM. Statistical comparisons were performed using the Mann-Whitney test. (ns=not significant). n=12.

**Figure EV3.**
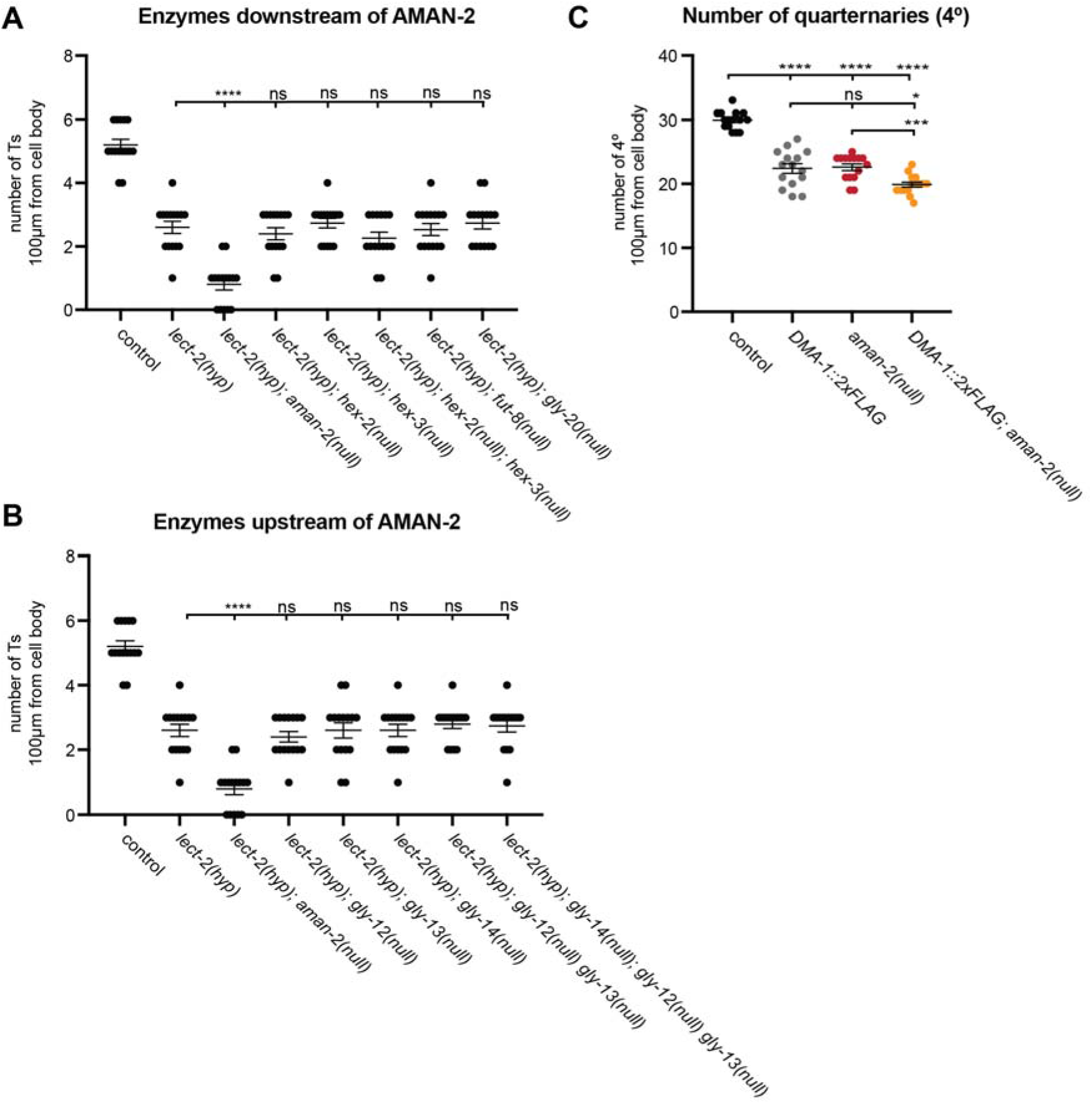
Quantification of branching in *N*-glycosylation pathway mutants. (A) Quantification of the number of “Ts” in wild type controls, *lect-2(gk864764*) hypomorphs, and *lect-2(gk864764*) in combination with mutants of MGAT1 orthologs. See strain list for alleles. Data are represented as mean ± SEM. Statistical significance was calculated using the Kruskal-Wallis test and is indicated (****p ≤ 0.0001; ns= not significant). n = 15 for all genotypes. (B) Quantification of the number of “Ts” in wild type controls, *lect-2(gk864764*) hypomorphs, and *lect-2(gk864764*) in combination with mutants of enzymes acting downstream of *aman-2*. See strain list for alleles. Data are represented as mean ± SEM. Statistical significance was calculated using the Kruskal-Wallis test and is indicated (****p ≤ 0.0001; ns= not significant). n = 15 for all genotypes. (C) Quantification of the number of quaternary branches in wild type control animals, DMA-1::2XFLAG (denoted as DMA-1 wildtype in Figure 4), and in combination with *aman-2*(*gk248486*). In the DMA-1::2XFLAG (*wy1041*) background, there is a baseline decrease in the number of branches, possibly due to the insertion of the tag (Dong *et al*., 2016). While equivalent to the phenotype of *aman-2*(*gk248486*), combining the two backgrounds results in a further decrease in quaternary branches, providing and additional example of how hypomorphic alleles in the Menorin pathway can be enhanced by loss *aman-2*. Data are represented as mean ± SEM. Statistical significance was calculated using the Kruskal-Wallis test and is indicated (*p ≤ 0.05, ***p ≤ 0.001, ****p ≤ 0.0001). n = 15 for all genotypes.

**Figure EV4.**
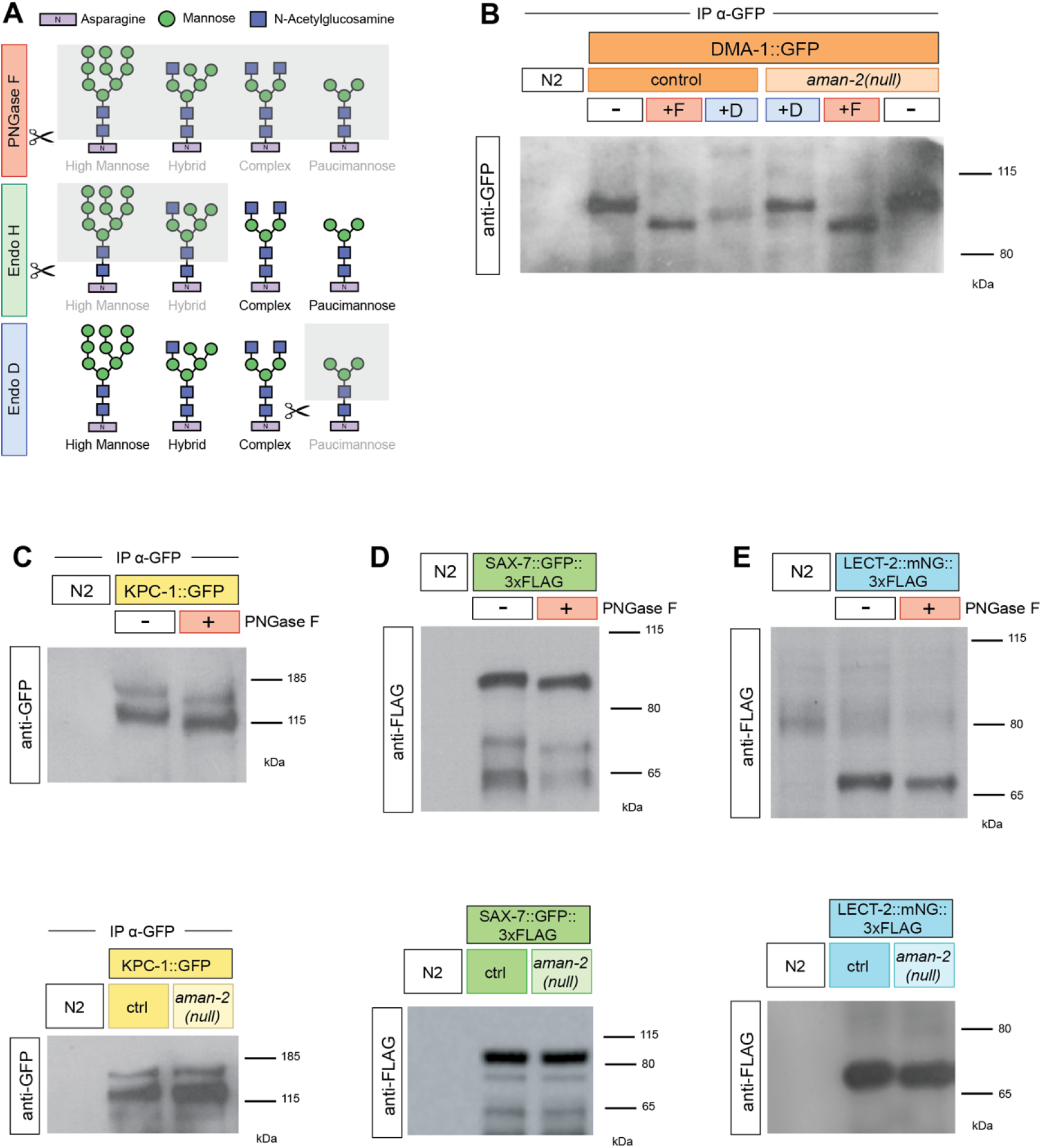
Members of the Menorin pathway are *N*-glycosylated. (A) Schematic of endoglycosidase activity on *N*-glycan chains. The gray shading represents parts of the chain that are cleaved by each respective endoglycosidase. Glycan residues are consistent with Figure 2A, and colors of endoglycosidases correspond to Figure 3E,G. (B) Western blot against GFP in *C. elegans* lysate DMA-1::GFP (*qyIs369*), after precipitating with anti-GFP antibody. Control indicates an otherwise wild type background as opposed to an *aman-2*(*gk248486*) background. The red boxed +F indicates that the lysate is treated with the PNGase F glycosidase, while the green boxed +D corresponds to the Endo D glycosidase, which cleaves paucimannose type *N*-glycans. Size shifts indicate that some paucimannose structures are present on DMA-1 (left), and that the *aman-2* mutant results in the loss of paucimannose structures on DMA-1 (right). Ladder is marked in kilodaltons (kDa). (C) Western blot against GFP in *C. elegans* lysate expressing no transgenes (N2) and expressing KPC-1::GFP (*dzEx1865*), after precipitating with anti-GFP antibody. The red boxed plus sign indicates that the lysate is treated with the PNGase F glycosidase. The downwards size shift reveals that *N*-glycan structures are present on KPC-1. In the bottom blot, control indicates an otherwise wild type background as opposed to an *aman-2*(*gk248486*) background. No size shift is observed. (D) Western blot against FLAG in *C. elegans* lysate expressing no transgenes (N2) and expressing SAX-7::GFP::3XFLAG (*dsIs290*). Robust expression precludes the need for immunoprecipitation. The red boxed plus sign indicates that the lysate is treated with the PNGase F glycosidase. The downwards size shift reveals that *N*-glycan structures are present on SAX-7. In the bottom blot, control indicates an otherwise wild type background as opposed to an *aman-2*(*gk248486*) background. No size shift is observed. The FLAG epitope contains no *N*-glycosylation sites. (E) Western blot against FLAG in *C. elegans* lysate expressing no transgenes (N2) and expressing endogenous LECT-2::mNeonGreen::3XFLAG (*dz249*). Robust expression precludes the need for immunoprecipitation. The red boxed plus sign indicates that the lysate is treated with the PNGase F glycosidase. The small downwards size shift reveals that *N*-glycan structures are present on LECT-2. In the bottom blot, control indicates an otherwise wild type background as opposed to an *aman-2*(*gk248486*) background. No size shift is observed.

## REFERENCES

Albeg A, Smith CJ, Chatzigeorgiou M, Feitelson DG, Hall DH, Schafer WR, Miller DM, 3rd, Treinin M (2011) C. elegans multi-dendritic sensory neurons: morphology and function. Mol Cell Neurosci 46: 308–317

Apweiler R, Hermjakob H, Sharon N (1999) On the frequency of protein glycosylation, as deduced from analysis of the SWISS-PROT database. Biochim Biophys Acta 1473: 4–8

Brenner S (1974) The genetics of Caenorhabditis elegans. Genetics 77: 71–94

Bülow HE, Hobert O (2006) The Molecular Diversity of Glycosaminoglycans Shapes Animal Development. Ann Rev Cell Dev Biol 22: 375–407

Chang IJ, He M, Lam CT (2018) Congenital disorders of glycosylation. Ann Transl Med 6: 477

Chen S, Spence Andrew M, Schachter H (2003) Isolation of null alleles of the Caenorhabditis elegans gly-12, gly-13 and gly-14 genes, all of which encode UDP-GlcNAc: α-3-D-mannoside β1,2-N-acetylglucosaminyltransferase I activity. Biochimie 85: 391–401

Chen S, Zhou S, Sarkar M, Spence AM, Schachter H (1999) Expression of three Caenorhabditis elegans N-acetylglucosaminyltransferase I genes during development. J Biol Chem 274: 288–297

Dennis JW, Nabi IR, Demetriou M (2009) Metabolism, cell surface organization, and disease. Cell 139: 1229–1241

Diaz-Balzac CA, Rahman M, Lazaro-Pena MI, Martin Hernandez LA, Salzberg Y, Aguirre-Chen C, Kaprielian Z, Bülow HE (2016) Muscle- and Skin-Derived Cues Jointly Orchestrate Patterning of Somatosensory Dendrites. Curr Biol 26: 2379–2387

Dickinson DJ, Goldstein B (2016) CRISPR-Based Methods for Caenorhabditis elegans Genome Engineering. Genetics 202: 885–901

Doitsidou M, Jarriault S, Poole RJ (2016) Next-Generation Sequencing-Based Approaches for Mutation Mapping and Identification in Caenorhabditis elegans. Genetics 204: 451–474

Dong X, Chiu H, Park YJ, Zou W, Zou Y, Ozkan E, Chang C, Shen K (2016) Precise regulation of the guidance receptor DMA-1 by KPC-1/Furin instructs dendritic branching decisions. Elife 5

Dong X, Liu OW, Howell AS, Shen K (2013) An extracellular adhesion molecule complex patterns dendritic branching and morphogenesis. Cell 155: 296–307

Dong X, Shen K, Bülow HE (2015) Intrinsic and extrinsic mechanisms of dendritic morphogenesis. Annu Rev Physiol 77: 271–300

Feng Z, Zhao Y, Li T, Nie W, Yang X, Wang X, Wu J, Liao J, Zou Y (2020) CATP-8/P5A ATPase Regulates ER Processing of the DMA-1 Receptor for Dendritic Branching. Cell Rep 32: 108101

Fogel AI, Li Y, Giza J, Wang Q, Lam TT, Modis Y, Biederer T (2010) N-glycosylation at the SynCAM (synaptic cell adhesion molecule) immunoglobulin interface modulates synaptic adhesion. J Biol Chem 285: 34864–34874

Freeze HH (2006) Genetic defects in the human glycome. Nat Rev Genet 7: 537–551

Gasser B, Saloheimo M, Rinas U, Dragosits M, Rodriguez-Carmona E, Baumann K, Giuliani M, Parrilli E, Branduardi P, Lang C et al (2008) Protein folding and conformational stress in microbial cells producing recombinant proteins: a host comparative overview. Microb Cell Fact 7: 11

Gilleard JS, Barry JD, Johnstone IL (1997) cis regulatory requirements for hypodermal cell-specific expression of the Caenorhabditis elegans cuticle collagen gene dpy-7. Mol Cell Biol 17: 2301–2311

Gupta R, Brunak S (2002) Prediction of glycosylation across the human proteome and the correlation to protein function. Pac Symp Biocomput: 310–322

Holt CE, Dickson BJ (2005) Sugar codes for axons? Neuron 46: 169–172

Inberg S, Meledin A, Kravtsov V, Iosilevskii Y, Oren-Suissa M, Podbilewicz B (2019) Lessons from Worm Dendritic Patterning. Annu Rev Neurosci 42: 365–383

Ioffe E, Stanley P (1994) Mice lacking N-acetylglucosaminyltransferase I activity die at mid-gestation, revealing an essential role for complex or hybrid N-linked carbohydrates.

Jaeken J, Peanne R (2017) What is new in CDG? J Inherit Metab Dis 40: 569–586

Jan YN, Jan LY (2010) Branching out: mechanisms of dendritic arborization. Nat Rev Neurosci 11: 316–328

Kaji H, Kamiie J, Kawakami H, Kido K, Yamauchi Y, Shinkawa T, Taoka M, Takahashi N, Isobe T (2007) Proteomics reveals N-linked glycoprotein diversity in Caenorhabditis elegans and suggests an atypical translocation mechanism for integral membrane proteins. Mol Cell Proteomics 6: 2100–2109

Kutscher LM, Shaham S (2014) Forward and reverse mutagenesis in C. elegans. WormBook: 1–26

Labasque M, Hivert B, Nogales-Gadea G, Querol L, Illa I, Faivre-Sarrailh C (2014) Specific contactin N-glycans are implicated in neurofascin binding and autoimmune targeting in peripheral neuropathies. J Biol Chem 289: 7907–7918

Lefebvre JL (2021) Molecular mechanisms that mediate dendrite morphogenesis. In: Current Topics in Developmental Biology, Academic Press:

Li L, Zinovyeva AY (2020) Protein Extract Preparation and Co-immunoprecipitation from Caenorhabditis elegans. J Vis Exp

Lu H, Wang SS, Wang WL, Zhang L, Zhao BY (2014) Effect of swainsonine in Oxytropis kansuensis on Golgi alpha-mannosidase II expression in the brain tissues of Sprague-Dawley rats. J Agric Food Chem 62: 7407–7412

Masu M (2016) Proteoglycans and axon guidance: a new relationship between old partners. J Neurochem 139 Suppl 2: 58–75

Medina-Cano D, Ucuncu E, Nguyen LS, Nicouleau M, Lipecka J, Bizot JC, Thiel C, Foulquier F, Lefort N, Faivre-Sarrailh C et al (2018) High N-glycan multiplicity is critical for neuronal adhesion and sensitizes the developing cerebellum to N-glycosylation defect. Elife 7

Metzler M, Gertz A, Sarkar M, Schachter H, Schrader JW, Marth JD (1994) Complex asparagine-linked oligosaccharides are required for morphogenic events during post-implantation development. EMBO J 13: 2056–2065

Minevich G, Park DS, Blankenberg D, Poole RJ, Hobert O (2012) CloudMap: a cloud-based pipeline for analysis of mutant genome sequences. Genetics 192: 1249–1269

Mire E, Hocine M, Bazellieres E, Jungas T, Davy A, Chauvet S, Mann F (2018) Developmental Upregulation of Ephrin-B1 Silences Sema3C/Neuropilin-1 Signaling during Post-crossing Navigation of Corpus Callosum Axons. Curr Biol 28: 1768–1782 e1764

Moloney DJ, Panin VM, Johnston SH, Chen J, Shao L, Wilson R, Wang Y, Stanley P, Irvine KD, Haltiwanger RS et al (2000) Fringe is a glycosyltransferase that modifies Notch [see comments]. Nature 406: 369–375

Moremen KW (2002) Golgi a-mannosidase II deficiency in vertebrate systems: implications for asparagine-linked oligosaccharide processing in mammals. Biochimica et Biophysica Acta 1573: 225–235

Ng BG, Freeze HH (2018) Perspectives on Glycosylation and Its Congenital Disorders. Trends Genet 34: 466–476

Okkema PG, Harrison SW, Plunger V, Aryana A, Fire A (1993) Sequence requirements for myosin gene expression and regulation in Caenorhabditis elegans. Genetics 135: 385–404

Oren-Suissa M, Hall DH, Treinin M, Shemer G, Podbilewicz B (2010) The fusogen EFF-1 controls sculpting of mechanosensory dendrites. Science 328: 1285–1288

Paschinger K, Hackl M, Gutternigg M, Kretschmer-Lubich D, Stemmer U, Jantsch V, Lochnit G, Wilson IB (2006) A deletion in the golgi alpha-mannosidase II gene of Caenorhabditis elegans results in unexpected non-wild-type N-glycan structures. J Biol Chem 281: 28265–28277

Paschinger K, Yan S, Wilson IBH (2019) N-glycomic Complexity in Anatomical Simplicity: Caenorhabditis elegans as a Non-model Nematode? Front Mol Biosci 6: 9

Poulain FE, Yost HJ (2015) Heparan sulfate proteoglycans: a sugar code for vertebrate development? Development 142: 3456–3467

Salzberg Y, Coleman AJ, Celestrin K, Cohen-Berkman M, Biederer T, Henis-Korenblit S, Bülow HE (2017) Reduced Insulin/Insulin-Like Growth Factor Receptor Signaling Mitigates Defective Dendrite Morphogenesis in Mutants of the ER Stress Sensor IRE-1. PLoS Genet 13: e1006579

Salzberg Y, Diaz-Balzac CA, Ramirez-Suarez NJ, Attreed M, Tecle E, Desbois M, Kaprielian Z, Bülow HE (2013) Skin-Derived Cues Control Arborization of Sensory Dendrites in Caenorhabditis elegans. Cell 155: 308–320

Salzberg Y, Ramirez-Suarez NJ, Bülow HE (2014) The proprotein convertase KPC-1/furin controls branching and self-avoidance of sensory dendrites in Caenorhabditis elegans. PLoS Genet 10: e1004657

Schroeder NE, Androwski RJ, Rashid A, Lee H, Lee J, Barr MM (2013) Dauer-specific dendrite arborization in C. elegans is regulated by KPC-1/Furin. Curr Biol 23: 1527–1535

Sekine SU, Haraguchi S, Chao K, Kato T, Luo L, Miura M, Chihara T (2013) Meigo governs dendrite targeting specificity by modulating ephrin level and N-glycosylation. Nat Neurosci 16: 683–691

Shah N, Kuntz DA, Rose DR (2008) Golgi alpha-mannosidase II cleaves two sugars sequentially in the same catalytic site. Proc Natl Acad Sci U S A 105: 9570–9575

Smith CJ, O’Brien T, Chatzigeorgiou M, Spencer WC, Feingold-Link E, Husson SJ, Hori S, Mitani S, Gottschalk A, Schafer WR et al (2013) Sensory Neuron Fates Are Distinguished by a Transcriptional Switch that Regulates Dendrite Branch Stabilization. Neuron 79: 266–280

Smith CJ, Watson JD, Spencer WC, O’Brien T, Cha B, Albeg A, Treinin M, Miller DM, 3rd (2010) Time-lapse imaging and cell-specific expression profiling reveal dynamic branching and molecular determinants of a multi-dendritic nociceptor in C. elegans. Dev Biol 345: 18–33

Stanley P, Taniguchi N, Aebi M (2015) N-Glycans. In: Essentials of Glycobiology, rd, Varki A., Cummings R.D., Esko J.D., Stanley P., Hart G.W., Aebi M., Darvill A.G., Kinoshita T., Packer N.H. et al (eds.) pp. 99–111. Cold Spring Harbor (NY)

Sundararajan L, Stern J, Miller DM, 3rd (2019) Mechanisms that regulate morphogenesis of a highly branched neuron in C. elegans. Dev Biol 451: 53–67

Tang LT, Diaz-Balzac CA, Rahman M, Ramirez-Suarez NJ, Salzberg Y, Lazaro-Pena MI, Bülow HE (2019) TIAM-1/GEF can shape somatosensory dendrites independently of its GEF activity by regulating F-actin localization. Elife 8

Thompson O, Edgley M, Strasbourger P, Flibotte S, Ewing B, Adair R, Au V, Chaudhry I, Fernando L, Hutter H et al (2013) The million mutation project: a new approach to genetics in Caenorhabditis elegans. Genome Res 23: 1749–1762

Tsalik EL, Niacaris T, Wenick AS, Pau K, Avery L, Hobert O (2003) LIM homeobox gene-dependent expression of biogenic amine receptors in restricted regions of the C. elegans nervous system. Dev Biol 263: 81–102

Vabulas RM, Raychaudhuri S, Hayer-Hartl M, Hartl FU (2010) Protein folding in the cytoplasm and the heat shock response. Cold Spring Harb Perspect Biol 2: a004390

Wei X, Howell AS, Dong X, Taylor CA, Cooper RC, Zhang J, Zou W, Sherwood DR, Shen K (2015) The unfolded protein response is required for dendrite morphogenesis. Elife 4: e06963

Zou W, Shen A, Dong X, Tugizova M, Xiang YK, Shen K (2016) A multi-protein receptorligand complex underlies combinatorial dendrite guidance choices in C. elegans. Elife 5

